# A new member in the Argonaute crew: the mt-miRNAs

**DOI:** 10.1101/2020.02.13.948554

**Authors:** Andrea Pozzi, Damian K. Dowling

## Abstract

Mutations within the mitochondrial genome have been linked to many diverse phenotypes. Moreover, the effects of these mutations have been shown to differ across sexes and environments. The mechanisms that explain the manifold array of mitochondrial genotypic effects on organismal function, and their context-dependency, have however remained a mystery. Here, we present evidence that mitochondria are involved in nuclear gene regulation via RNA interference; transcribing mitochondrial (mt-)miRNAs that may repress the transcription of nuclear genes that previously had no known involvement in mitochondrial function. Our findings uncover a new mechanism by which mitochondria may shape the expression of animal life-histories and health components; implying that the influence of the mitochondria in regulating organismal function extends well beyond the process of energy production.

## Introduction

Interest in mitochondrial biology is on the rise, with a growing number of studies highlighting the complex role of the mitochondria in cell regulation (Picard, Wallace, and Burelle 2016; Sloan et al. 2018; Sprenger and Langer 2019). Among these, numerous studies have found that sequence variation in the mitochondrial DNA (mtDNA) can affect the expression of a range of life-history and health related traits, from fertility, to longevity and thermal tolerance (Lajbner et al. 2018; Camus et al. 2017; Rand, Fry, and Sheldahl 2006; Song and Lewis 2008; Yee, Sutton, and Dowling 2013; James and Ballard 2003). Furthermore, while it is well known that the mtDNA can harbour loss-of-function mutations conferring mitochondrial disease in humans (Wallace 2018), emerging studies implicate mtDNA mutations in a range of other late-onset diseases not previously linked to mitochondrial function (Hudson et al. 2014). For example, Hopkins et al. (2017) recently reported an association between the frequency and type of mtDNA mutations and aggressiveness of prostate cancer. Furthermore, the phenotypic effects of these mtDNA mutations appear to be routinely moderated by the nuclear genetic background alongside which the mtDNA mutations are co-expressed (Hill et al. 2018), suggesting a broad role for intergenomic regulation (“mitonuclear communication”) involving exchange of proteins, metabolites and genetic products between genomes (Wu et al. 2019; Moriyama, Koshiba, and Ichinohe 2019; Zhu, Ingelmo, and Rand 2014). The mechanistic basis of the molecular interactions that underpin mitonuclear regulation of cellular and organismal function, however, remains elusive.

In 2019, Kopinski et al. provided new insights into how the mitochondria communicate with the nucleus, reporting a key role for mitochondrial metabolites and subcellular redox levels. The authors found that variation in the level of intracellular mtDNA heteroplasmy (i.e. the frequency of normal to mutant mtDNA molecules) modulates mitochondrial metabolites, influencing the abundance of substrate necessary for methylation and acetylation of specific histones, thus affecting patterns of nuclear expression (Kopinski et al. 2019). Their findings were to some degree consistent with those of a previous study by Guantes et al. (2015), who reported strong correlations between mtDNA content and changes to the epigenetic and transcriptional profile of the cell. However, Guantes et al. (2015) found that some of the mtDNA-content mediated changes to cellular regulation were unlikely related to the presence of mitochondrial metabolites. Furthermore, they found that mtDNA content also shapes patterns of post-transcriptional regulation; a type of regulation that is not usually affected directly by histone methylation. Mitochondrial-mediated modification to levels of protein translation, affected by post-transcriptional modifications to gene expression, would require a mechanism that is able to interfere with mRNA translation. To date, such a mechanism is unknown to exist.

Mitochondrial RNAs (mt-RNAs) represent a possible mediator of patterns of mitochondrial-mediated post-transcriptional regulation. Functional mt-RNAs are well-known in the form of the 22 tRNAs and 2 rRNAs encoded by the typical bilaterian mitochondrial genome. However, novel types of RNAs of mitochondrial origin have recently been identified. For example, Dhir et al. (2018) described a new class of double-stranded RNAs encoded in the mitochondria that are able to trigger antiviral signaling in humans (Dhir et al. 2018). Although to date these double-stranded RNAs have only been identified in humans, different types of novel small mitochondrial RNAs have been described in multiple species across two metazoan phyla, Chordata and Mollusca (Ro et al. 2013; Riggs et al. 2018; Larriba, Rial, and Del Mazo 2018; Bottje et al. 2017; Mercer et al. 2011; Pozzi et al. 2017; Pozzi and Dowling 2019). Yet, despite increasing interest in the putative role these small mitochondrial RNAs may play in the regulation of cellular function, clear evidence of their functionality remains absent.

Given similarities in their length and sequence to microRNAs, Pozzi et al. (2017) hypothesized involvement of small mitochondrial RNAs in the regulation of mRNA translation through RNA interference (RNAi) (Pozzi et al. 2017). RNAi is a process in which a microRNA (miRNA) leads a protein complex to block translation of a target mRNA (Ambros 2004; Ha and Kim 2014). miRNAs are partially complementary to a regulatory region of the mRNAs, and due this close miRNA-mRNA affinity, the protein complex is able to precisely bind its target mRNA and hinder its binding to the ribosome (Ambros 2004; Cloonan 2015). Within this protein complex, the main protein binding the miRNAs is Ago2, an endonuclease shared across multiple species necessary for RNAi (Ha and Kim 2014; Cloonan 2015). Interestingly, Ago2 has previously been reported to co-localize with mitochondria (Bandiera et al. 2011; Zhang et al. 2014) and, moreover, to associate with mitochondrial tRNA (mt-tRNA_Met_) in the cytoplasm (Maniataki and Mourelatos 2005). Accordingly, Ago2 is an excellent candidate to further probe the hypothesis that the small mitochondrial RNAs serve a similar role as nuclear encoded miRNAs in RNAi, and therefore constitute mitochondrial miRNAs (hereafter, “mt-miRNAs”).

Here, we investigate mitochondrial involvement in RNAi by verifying the presence of multiple features of miRNAs in the mitochondrial small RNAs. Firstly, we sought to verify binding between the mitochondrial small RNAs and Ago2. To this end, we leveraged published datasets of RNA sequencing (RNA-seq) and RNA-binding-protein co-immunoprecipitation sequencing (RIP-seq) (Townley-Tilson, Pendergrass, and Marzluff 2006). These datasets come from pre-published studies that reported novel mechanistic insights into RNAi. We re-purposed these datasets to investigate the capacity for mtDNA-mediated involvement in RNAi. Secondly, we tested whether the mitochondrial small RNAs are generated by pre-mitochondrial small RNAs, similarly to the case of miRNAs. To achieve this, we screened for the presence of matching transcriptional profiles between datasets of long and small RNAs extracted from the same individuals, which enabled us to determine if the mt-miRNAs are transcribed from transcripts of ~70nt length, as commonly happens in the miRNAs (Ha and Kim 2014). Thirdly, given that miRNAs have been shown to be conserved across multiple clades, we explored levels of sequence conservation in the mitochondrial small RNAs. We compared the presence of mitochondrial small RNAs that exhibit features like the miRNAs across multiple model organisms by using small RNAs-seq datasets from multiple independent and taxonomically diverse studies. Finally, we investigated the presence of mRNA targets for the most conserved of the small mitochondrial RNAs. We used a mix of computational and experimental methods to determine the presence of a target for the mt-miRNA. By combining multiple datasets and approaches, our study provides the first evidence of a functional relationship between the mitochondrial small RNAs and RNAi; thus, supporting the hypothesis that these RNAs are mitochondrially transcribed miRNAs (mt-miRNAs).

## Results

### Mt-miRNAs bind Ago2

To investigate the ability of the small mitochondrial RNAs to bind Ago2, we analysed a RIP-seq dataset including two different human cell lines; neural progenitor and teratoma-derived fibroblast. This dataset includes all the small RNAs that bind to Ago2, however, our focus was only on the mt-miRNAs, which have been previously ignored. Accordingly, we identified mt-miRNAs binding Ago2 in each of the two cell lines (**Fig.1**). Through this analysis, we found that the mt-miRNAs are present only in the Ago2 immunoprecipitations (IPs), mostly in the regions coding for mt-tRNAs, while the control IP samples have almost no mt-miRNA present. The expression of the mt-miRNAs differs across the different cell line types, which are reflective of different tissues, confirming previous studies demonstrating that mitochondrial transcription is usually tissue-specific (Mercer et al. 2011; Pozzi and Dowling 2019; Scheibye-Alsing et al. 2007). Aside from Ago2 RIP-seq, the authors of the original dataset performed other treatments on their samples to verify genuine binding of their focal miRNAs with Ago2. One of these treatments is particularly significant for our study: RNase I treatment. RNase I is an endonuclease able to digest RNAs that remain unbound to proteins, thus higher concentrations of RNAase I will be more effective in eliminating RNA contamination. We analysed samples treated with different RNase I concentrations and found that higher concentrations of RNase I have no effect on transcription levels of mt-miRNAs (See SI Fig.S1). Furthermore, we performed a gene-by-gene analysis on both Ago2-IP and control IP samples to verify whether the transcriptional signatures of the mt-miRNAs are consistent with them representing functional RNAs or noise (Pozzi et al. 2017; Mercer et al. 2011). These analyses showed that the mt-miRNAs with the highest level of transcription are encoded within mt-RNAs, confirming what found by previous studies in other human cell lines (Mercer et al. 2011; Ro et al. 2013). Due to the large number of genes analysed, the results of the gene-by-gene analysis are in the SI Fig.S2. Our analysis of the Ago2-IP samples provides the first evidence that the mt-miRNAs bind to Ago2, demonstrating their involvement in RNAi.

**Fig 1.**
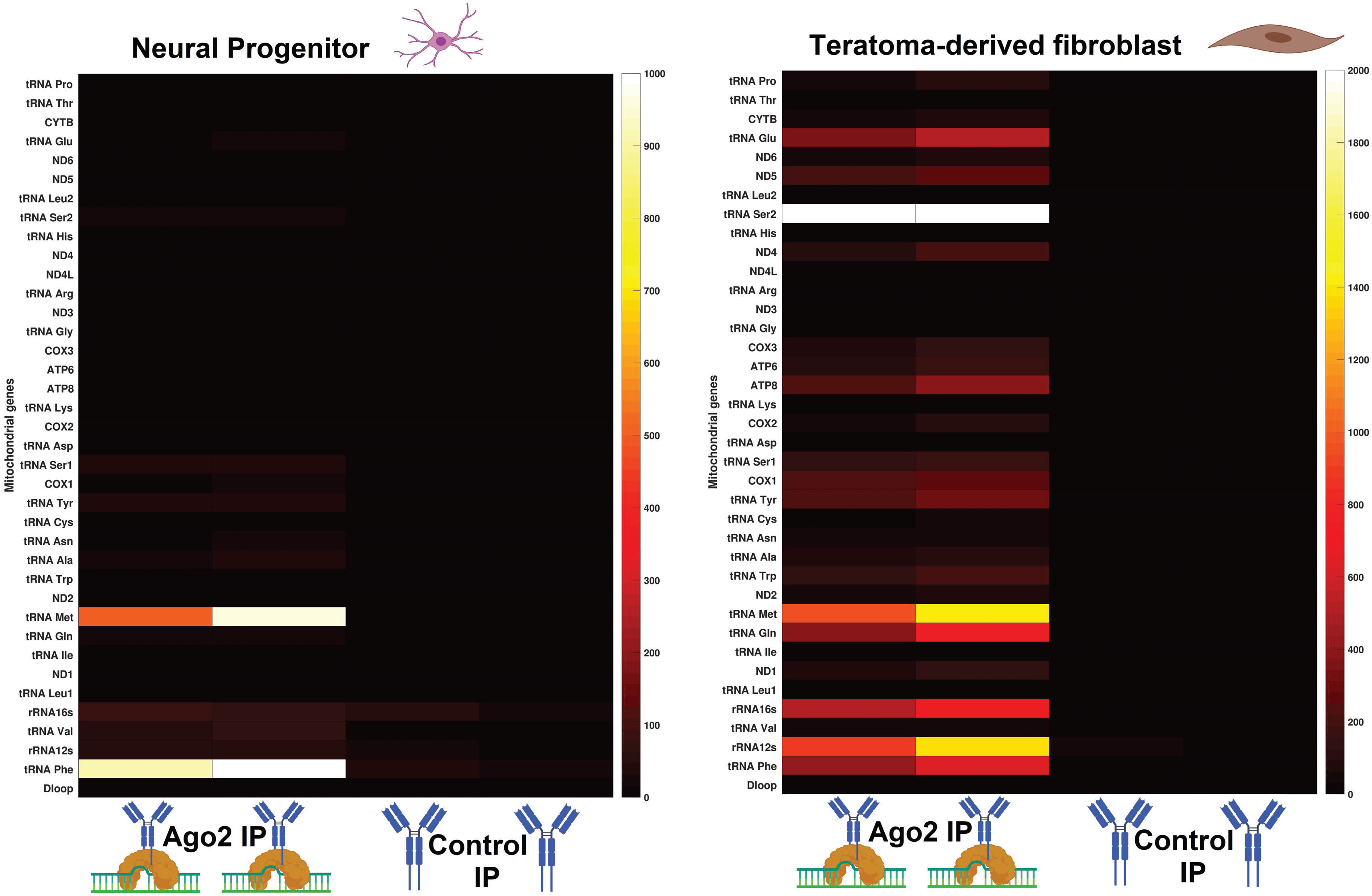
The mt-miRNAs bind Ago2. In two different cell lines, Ago2-IP samples are enriched in expression of specific mt-miRNAs when compared to a control IP, which contains only the IgG antibody. On the Y-axis, all the canonical mitochondrial genes are listed. On the X-axis, there are two biological replicates for each treatment. The two heatmaps are divided because they represent experiments from different cell lines. The sample type and treatments are illustrated through BioRender icons. The color scale is black-red-white, in which black and white indicate the lowest and highest levels of expression respectively. As the mt-miRNAs are an uncharacterized type of RNAs, we did not perform any normalization, thus the expression is measured in raw read counts. Gene-by-gene analyses of the transcriptional signatures are shown in Figure S2.

### Mt-miRNAs are encoded in mt-tRNAs and protein-coding genes

To further investigate the transcriptional signature of mt-miRNAs across tissues and species, we expanded the number of datasets analysed, by including more cell lines from humans and mice from other independent studies. This analysis confirmed the presence of mt-miRNAs binding to Ago2 in human (HeLa) and mouse embryonic cell lines (Scherer, Syverton, and Gey 1953) (**Fig.2**). To verify the enrichment of mt-miRNAs in Ago2-IP samples, we calculated the fold change between samples of the same cell line in which one samples had undergone the IP process, and the other had not. Through this experiment, we identified an enrichment in mt-miRNAs across Ago2 IP samples in both species. In HeLa cells, a mix of tRNAs and protein-coding genes are enriched for these mt-miRNAs, providing the first evidence the mt-miRNAs can be encoded within both mt-tRNAs and protein coding genes. Nonetheless, the most enriched mt-miRNA (5fold higher expression) is encoded in the mt-tRNA Met. This finding supports a previous study which found that mt-tRNA Met binds to Ago2 outside the mitochondria (Maniataki and Mourelatos 2005). The high expression of the mt-miRNA_Met_ in these samples (HeLa cells), and in the previous analysis (neural progenitor and teratoma-derived fibroblast cell lines, Fig 1) suggests that although the mt-miRNAs have tissue-specific expression, some mt-miRNAs are conserved across tissues. The analysis of the mouse samples provides similar results. We found the mouse embryonic stem cells are enriched for mt-miRNAs encoded across multiple genes. Notably, the genes enriched for mt-miRNAs in the mouse samples differed from those in the human HeLa cell lines, with mt-ATP8 the only mt-miRNA exhibiting high expression in the Ago2-IP samples of both species.

**Fig 2.**
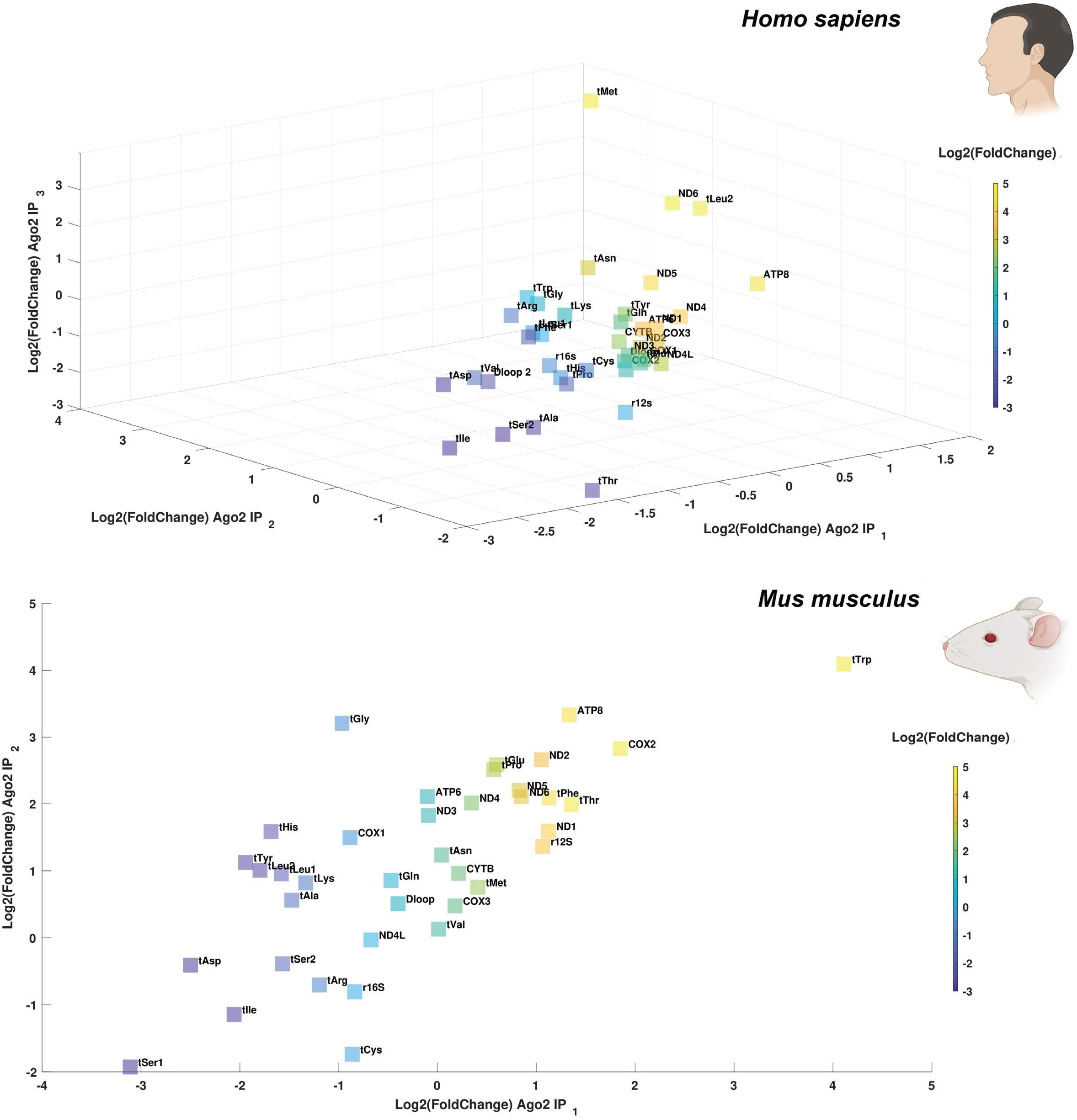
The mt-miRNAs are enriched in Ago2-IP samples. The plots show that Ago2-IP samples are enriched in expression for specific mt-miRNAs when compared to the same sample without Ago2-IP treatment. The enrichment is calculated though Log2 fold change, thus a fold change of two means that the gene has four times higher expression when compared to its counterpart without Ago2-IP treatment. We used a color scale of blue-green-yellow, in which blue and yellow indicate the lowest and highest levels of expression respectively. As the mt-miRNAs are an uncharacterized type of RNAs, we did not perform any normalization, thus the expression is measured in raw read counts. The species used in each plot is indicated through small BioRender icons. **A)** The 3D-plot shows patterns of mt-miRNA expression in Ago2-IP treated samples of HeLa cells. The three axes represent three different biological replicates for each of the two treatments (IP and non-IP). **B)** The plot shows patterns of mt-miRNA expression the Ago2-IP treated samples derived from mouse embryonic stem cells. The two we represent two different biological replicates for each of the two treatments (IP and non-IP). Gene-by-gene analyses of the transcriptional signatures are shown in Figure S3 (human) and Figure S4 (mouse).

To better understand the differences and similarities in the results between human and mouse samples, we performed a gene-by-by gene analysis of the transcriptional signature of the mt-miRNAs across the mouse embryonic stem cells and HeLa cell lines (See SI Fig.S3-4). Surprisingly, closer scrutiny of the putative mt-miRNA at mt-ATP8 in both species showed that the mt-miRNA is only likely to exist in humans (**Fig.3A**). In fact, while the human mt-ATP8 encodes an mt-miRNA (32nt long) with clear start and end, the same gene in mouse has only noise, without any clear transcriptional signature presence. This suggests that the enrichment patterns of the mt-miRNAs are accurate in predicting the presence of mt-miRNAs only when paired with detailed analysis of the mt-miRNAs transcriptional signature. Furthermore, we identified unusual transcriptional signatures that might shed light on some aspects of the biogenesis of the mt-miRNAs. Indeed, some mt-miRNAs are not fully encoded within a gene but overlap across two different genes (**Fig.3B**). We were able to identify this phenomenon only in the mouse samples, where both mt-miRNA Phe and mt-miRNA Thr partially overlap with the neighbour gene (an overlap of up to 6bp). Moreover, both of these mt-miRNAs have isoforms, “mt-isomiRs”, similar to observations in previous studies of nuclear miRNAs (Budak et al. 2016; Tan et al. 2014; Desvignes et al. 2015). These mt-miRNAs feature two different isoforms of different lengths: the short mt-isomiR ends where the first gene ends, while the longer mt-isomiR overlaps on the second gene by several nucleotides. Interestingly, these isoforms are quite long compared to other small RNAs such as miRNA (Ha and Kim 2014). In fact, the mt-isomiRs Phe are 32nt and 37nt long respectively, while the mt-isomiRs Thr are 37nt and 43nt long. Due to the length of these mt-isomiRs, it is possible that the longer isoform may represent a transitional stage for the shorter mature form. This phenomenon would be similar to what happens in piwi-interacting RNAs (piRNAs), in which proteins located on the surface of the mitochondria edit the length of piRNAs during their maturation process (Kim 2006; Nishimura et al. 2018; Ding et al. 2017; Bronkhorst and Ketting 2018). While the mechanism underpinning the generation of these mt-isomiRs remains unknown, our analysis provides the first report of mt-miRNAs encoded across the boundaries of genes, and, in general, the presence of mitochondrial products encoded across genes.

**Fig 3.**
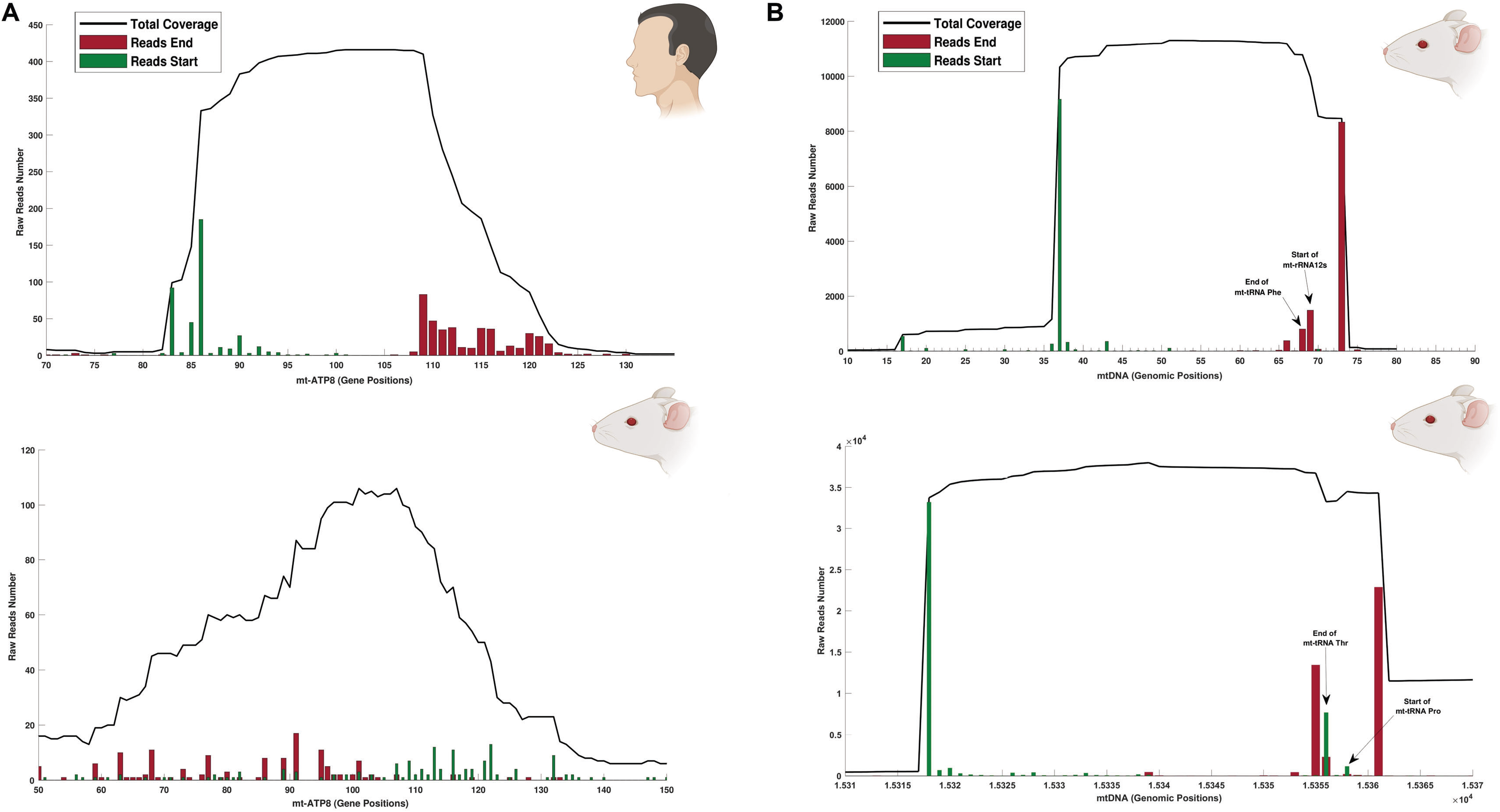
mt-miRNAs are encoded across genes. A) shows the coverage of small RNAs within the gene mt-ATP8 in human HeLa cell lines (upper left hand panel) and mice embryonic stem cells (lower left hand panel), and highlights the difference between a transcriptional profile showing a genuine mt-miRNA (human), and noise from the alignment of random small mitochondrial RNAs (mouse). The total coverage for each position in the gene is indicated with a black line, while the number of reads starting and ending at each position is indicated using green and red bars respectively. This type of representation provides the resolution necessary to verify the presence or absence of an mt-miRNA within a mitochondrial gene. As the mt-miRNAs are an uncharacterized type of RNAs, we did not perform any normalization, thus the expression shown on the Y-axis is measured in raw read counts. The X-axis shows the gene positions of each read. **B)** The panels represented on the right side of the figure highlight the transcriptional profile of two mt-miRNAs encoded across genes in mice embryonic stem cells. The first mt-miRNA (upper right hand panel) is encoded mostly within the mt-tRNA Phenylalanine; however, a long isoform of this mt-miRNA includes five nucleotides of the gene mt-rRNA12s. The second mt-miRNA (lower right hand panel) is encoded mostly within the mt-tRNA Threonine; however, a long isoform of this mt-miRNA includes six nucleotides.of the gene mt-tRNA Proline. Small black arrows indicate either the end or start of mt-genes where relevant.

### Mt-miRNAs and nuclear miRNAs have a different maturation process

Due to the presence of mt-isomiRs of different length, we investigated the possibility of these RNAs representing transition stages of the mt-miRNA maturation process. To address this, we re-purposed an RNA-seq dataset used for a study of the renal disease, Autosomal polycystic kidney disease, which includes several kidney samples from transgenic mice (Woo et al. 2017). The analysed dataset contains RNA samples in which the researchers had extracted both small and long RNAs, separately from the same individual mice (Woo et al. 2017). By comparing the transcriptional signature of small and long RNAs in the same samples, we can screen for the presence of matching transcriptional signatures, to determine whether either the start or end of each mt-miRNA matches with the expression of a mt-long RNA. However, we did not identify any matching transcriptional signatures between small and long RNAs (**Fig.4** SI Fig.S5). The small RNA samples show high levels of expression of mt-miRNAs in the mt-tRNAs Met and Ser 1, while the long RNAs samples have no distinct transcriptional hotspots. In fact, although most biological replicates of the long RNAs samples have a similar pattern of expression, we were not able to identify any signal suggesting the presence of functional RNAs. Thus, we argue that the consistent noisy pattern present in the long RNAs samples is most likely explained by the different chemical properties of the sequences, which would lead to some parts of the sequence being slightly overrepresented than others (Ross et al. 2013). On the contrary, the small RNA dataset exhibited a pattern concordant with the expression of mt-miRNAs: ~30nt sequences with clear cut start and end positions, consistent across multiple biological replicates (SI Fig S5). Thus our results suggest that the mt-miRNAs are not processed from pre-mt-miRNAs, as previously suggested, but are more likely to be matured directly from the polycistronic mt-RNA (Mercer et al. 2011; Rorbach and Minczuk 2012) from which mt-tRNAs and mt-rRNAs are matured.

**Fig 4.**
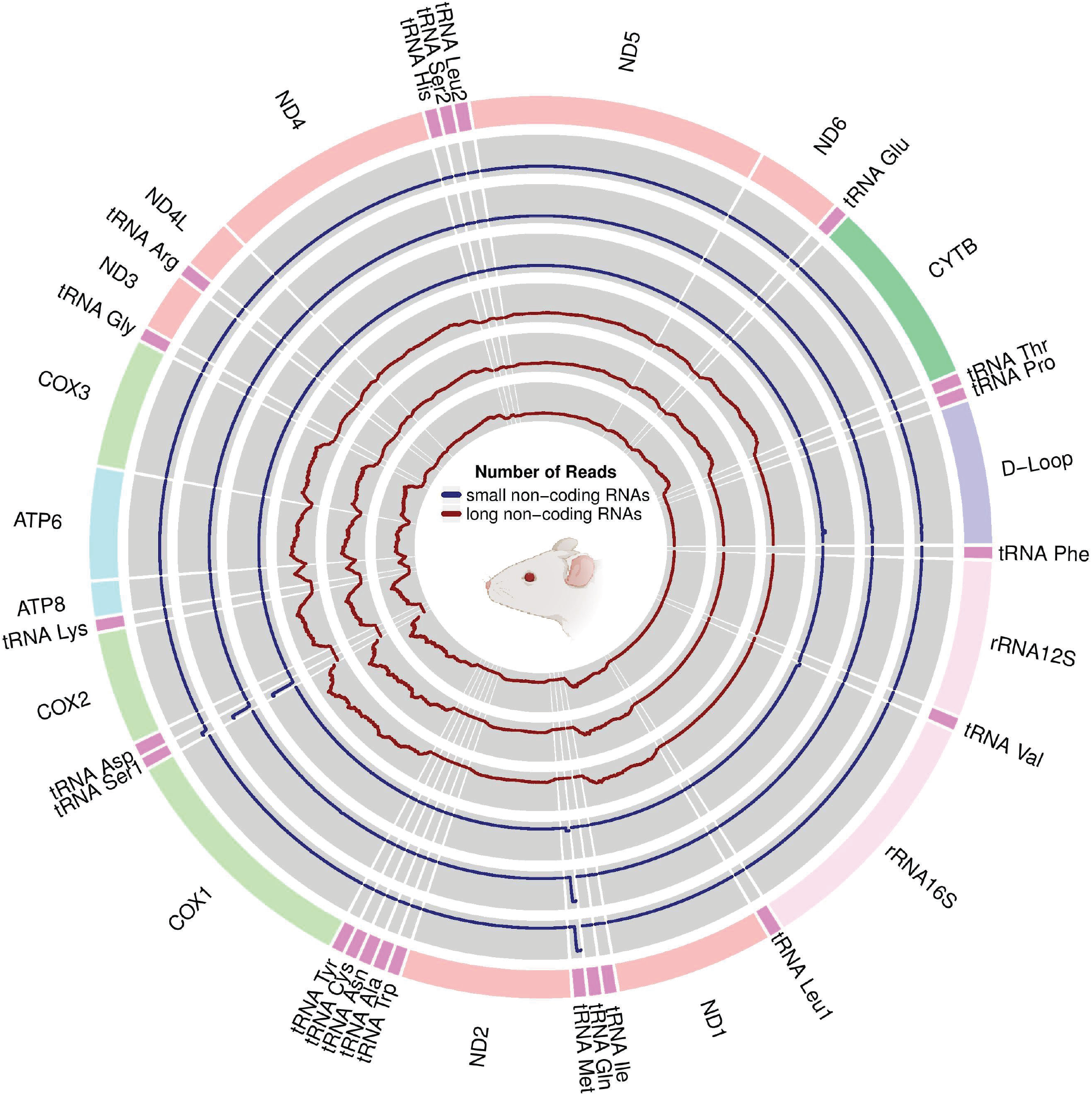
mt-miRNAs are not matured through pre-mt-miRNAs. The figure represents the transcriptional profile of all canonical mitochondrial genes for both small and long RNAs. The expression of small RNAs in each gene is indicated in dark blue, while the expression of long non-coding RNAs, both polyadenylated and not, is represented in dark red. As the mt-miRNAs are an uncharacterized type of RNAs, we did not perform any normalization, thus the expression is measured in raw read counts. To see more replicates, check Figure S5.

### mt-miRNA_Met_ is conserved across *Chordata*

Nuclear miRNAs are usually conserved across species (Kenny et al. 2015; Lee, Risom, and Strauss 2007), and we investigated whether mt-miRNA_Met_, which is conserved in both human and mouse (Fig. 1, 4), would be conserved in other model organisms. Thus, we analysed the expression and primary sequence of mt-miRNA_Met_ across two tissues of four model organisms and verified its conservation in *Chordata* (**Fig 5**). The four model organisms (human, mouse, chicken, and zebrafish) show a very consistent transcriptional signature for the mt-miRNA_Met_ across both brain and liver samples (**Fig 5A**). In fact, in all species the mt-miRNA_Met_ 3’ end is at the beginning of the mt-tRNA Met gene, while the 5’ end is ~32nt after. However, one of the tissues has a different signature. Samples from the human liver exhibit a noisier transcriptional signature, suggesting that despite its broad conservation, this mt-miRNA might not be ubiquitously expressed across all tissues. Indeed, this result aligns with one of our previous findings in mouse, as we did not find the mt-miRNA_Met_ in mouse embryonic stem cells, either transcribed or bound to Ago2. Similarly, to the transcriptional signature, we found that the primary sequence of the mt-miRNA_Met_ was highly conserved across the four model organisms (**Fig 5B**). The mt-miRNA_Met_ sequence is almost identical in all species, having only one polymorphism in amniotes, and four in zebrafish. When comparing the number of polymorphisms in the mt-miRNA_Met_ to the polymorphisms in the rest of the mt-tRNA Met, we found that the polymorphisms are underrepresented in the mt-miRNA_Met_. In fact, in human, mouse and chicken there are around 4 times more mutations in the rest of the mt-tRNAMet compared to the mt-miRNA_Met_ (1/30 against 5/39), while in zebrafish there are around 2 times more (4/30 against 10/39). This suggests that the region harbouring the mt-miRNA_Met_ might be under stronger purifying selection than its counterpart, which might be explained by the presence of overlapping selection due to the dual role of these regions in encoding both mt-miRNAMet and mt-tRNA Met.

**Fig 5.**
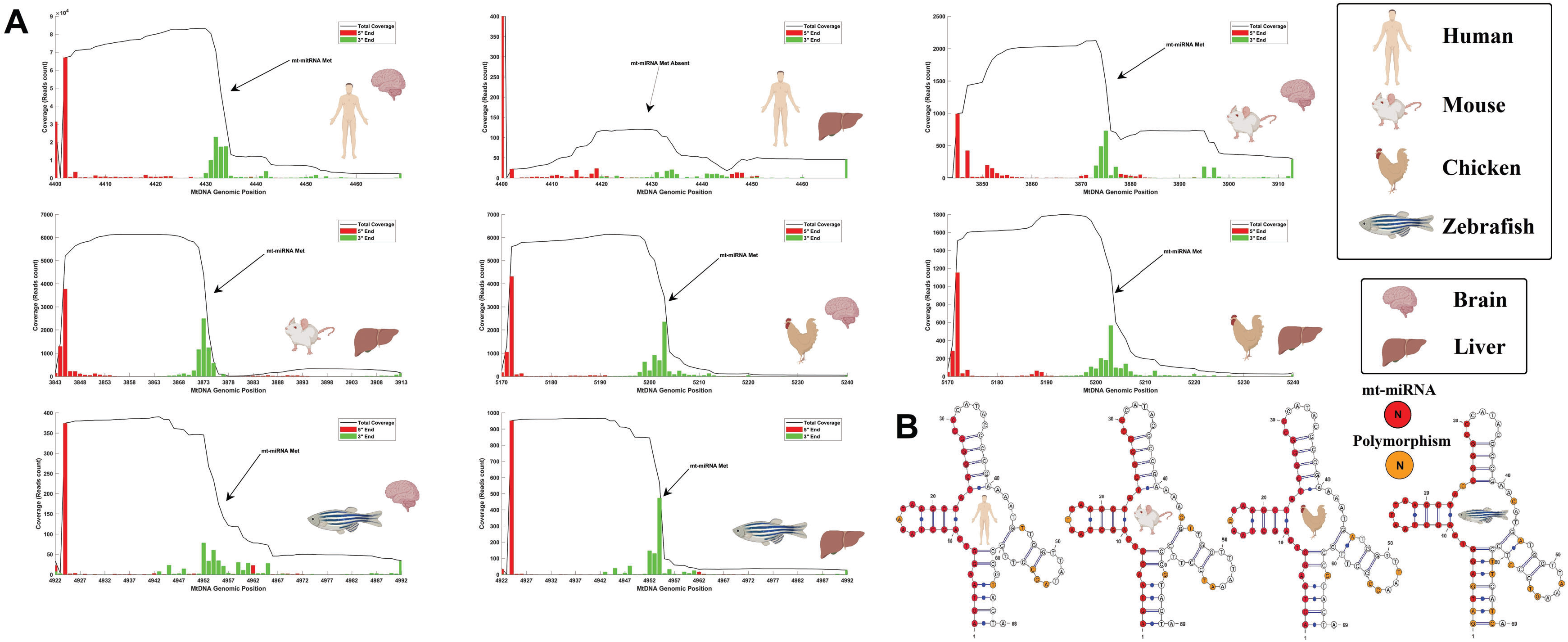
mt-miRNA_Met_ is conserved across species. In the figure panels, the expression and sequence of the mt-miRNA_Met_ are represented for several major model organisms: human (*Homo sapiens*), mouse (*Mus musculus*), chicken (*Gallus gallus*), and zebrafish (*Danio rerio*). For each species, tissues from two different organs are considered; brain and liver. The species and tissue of each sample are indicated through BioRender icons. **A**) The coverage (read count) in the mitochondrial genome, corresponding to the region of the mt-tRNA Met and mt-miRNAMet, is depicted. With exception of human liver tissue, the mt-miRNA Met is present across all species and tissues. The Y-axis represents the expression in raw reads count, while the X-axis represents the genomic positions in the mitochondrial genome of reference for each species. **B**) We represented the mt-tRNA Met of each species while highlighting in red the sequences corresponding to the mt-miRNA_Met_. We highlight in orange the presence of polymorphisms across the species, by comparing reference mitochondrial genomes of each species.

### The mt-miRNA_Met_ targets CFLAR in human temporal lobe

We investigated the function of the very conserved mt-miRNA, mt-miRNA_Met_, by verifying the presence of target mRNAs in the most well-characterized species, humans. To investigate the targets of mt-miRNA_Met_, we screened all 67087 human transcripts using a computational target predictor, which found 8709 potential targets (**Fig 6A**). These targets have different scores, as some of them are more likely to be a genuine target compared to the others. However, since it is impossible to establish a fully objective threshold, we filtered these potential targets for a specific function. Although probably this mt-miRNA target many transcripts, we decided to focus on genes involved in insulin regulation, because this pathway is at the intersection of many diseases in which mitochondrial mutations and nuclear miRNAs appear to be involved (C. Lee et al. 2015; Mohlke et al. 2005; Chalkia et al. 2018; Duarte, Palmeira, and Rolo 2015; Heni et al. 2015). By filtering for insulin regulation, we identified 74 transcripts. These transcripts were then validated by using brain Ago2-IP samples enriched in miRNAs targets (**Fig 6B**). These brain samples are an ideal tissue to study the presence of these targets, given that mitochondrial mutations are related to many neurodegenerative diseases (Takasaki 2009; Dölle et al. 2016). Through this analysis we were able to validate one of the transcripts of the gene CASP8 and FADD-like apoptosis regulator (CFLAR), which, as the name suggests, is a key protein in the regulation of a caspase (CASP8) involved in the apoptosis pathway. However, as the mt-miRNA_Met_ might not be the only miRNA binding this region, we verified how many nuclear miRNAs are able to bind the same region. By using programs for *in silico* prediction of miRNA targets in humans, we found that 13 nuclear miRNAs can bind the same region (**Fig 6C**). Nonetheless, by analysing other Ago2-IP samples from a similar part of the brain we found that none of these nuclear miRNAs are expressed in brain tissue, while the mt-miRNA_Met_ is present (**Fig 6D**). To verify that the absence of these miRNAs was not due to technical mistakes, or bias of the library, we analysed three other common miRNAs (let-7, mir9, and mir100) finding that they are expressed in the brain, and in some instances, at a similar expression level to the mt-miRNAMet. This analysis thus confirms that mt-miRNA_Met_ binds to the CFLAR UTR.

**Fig 6.**
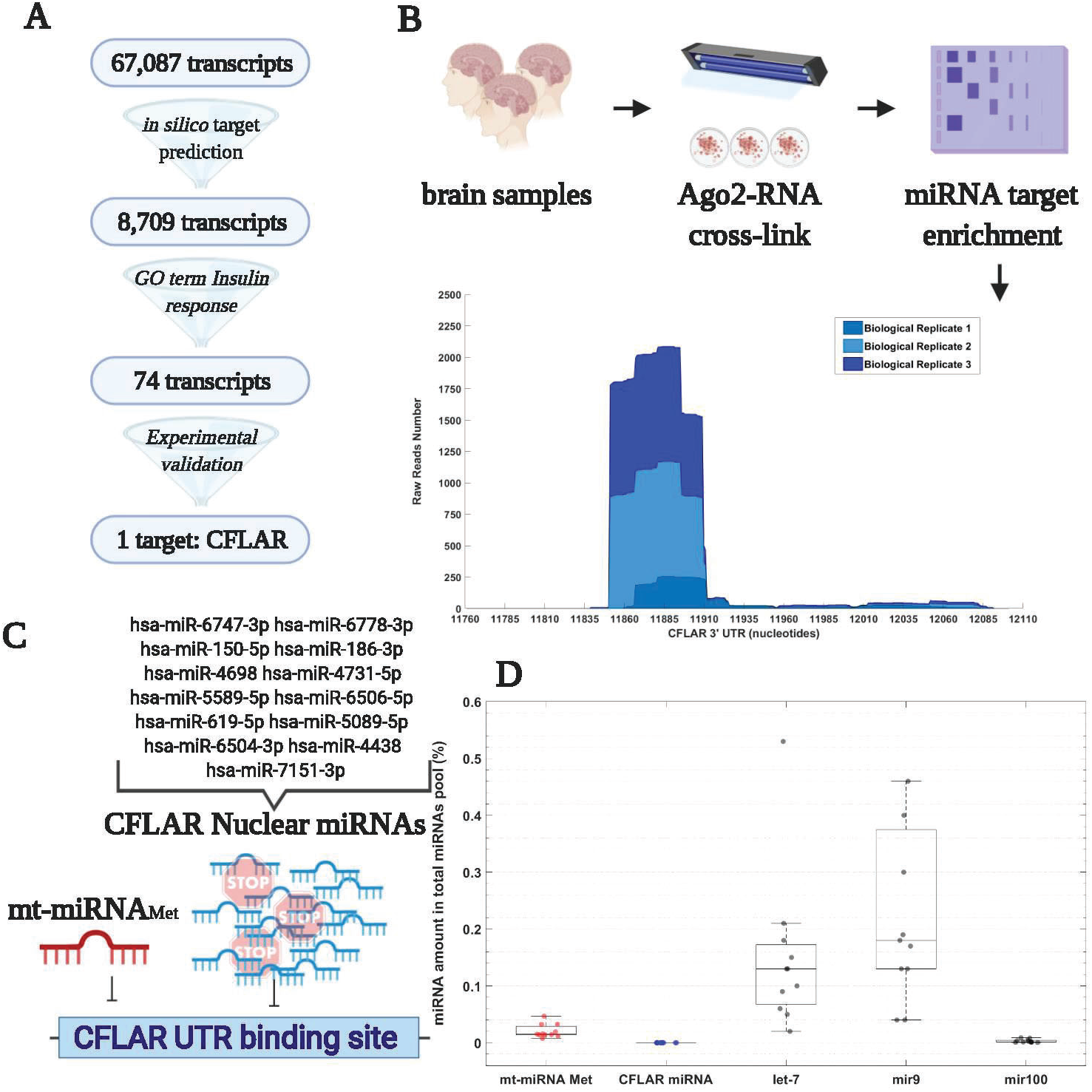
mt-miRNA_Met_ targets the gene CFLAR in brain tissue. The figure represents the identification of an mRNA target for the mt-miRNA_Met_. **A**) A schematic representation of the target prediction used to identify the CFLAR gene. Briefly, we analyzed the UTR of all human transcripts, identifying 8,709 possible targets through *in silico* prediction. Across these targets, we extracted only the ones involved in insulin response, a mechanism key in many mitochondria-related diseases. We then experimentally validated these targets by using an Ago2-IP dataset. **B**) We experimentally validated the binding between Ago2 and CFLAR by analyzing a dataset composed of three independent brain tissue samples. In this dataset Ago2 has been cross-linked to target mRNAs, and these RNAs have been isolated by size selection in gel. We show the high coverage present in the region predicted to bind mt-miRNA_Met_ on the CFLAR 3’ UTR across three biological replicates. The X- and Y- axes represent the positions within the 3’ UTR of CFLAR and the abundance of reads. **C**) a schematic representation of the binding of mt-miRNA_Met_ and the CFLAR nuclear miRNAs, a group of miRNAs predicted to bind this region of CFLAR in humans. A list of these miRNAs is present on the side of the figure, using the nomenclature present on miRBase. **D**) The boxplot represents the expression of four miRNAs, and a group of miRNAs named here CFLAR miRNAs. The expression is represented as the percentage representation of a given miRNA relative to the entire pool of miRNAs. Due to the absence of all the CFLAR miRNAs, we represented them as one group of pooled miRNAs (CFLAR miRNA). The presence of let-7, mir9, and mi100 to highlight that common miRNAs are expressed in a comparable way to mt-miRNA_Met_ in the brain samples that we analyzed.

To better understand the function of mt-miRNA_Met_, we investigated the function and evolution of its target, CFLAR. By comparing the CFLAR genomic region across six primates with high-level nuclear genome sequencing, we found that the human CFLAR has a unique structure that might be related to the presence of the binding site of mt-miRNA_Met_ (**Fig7A**). Indeed, the binding site of mt-miRNA_Met_ is near the end of the 12kb 3’ UTR of CFLAR. This is puzzling given the human CFLAR transcript consists of ~2kb of protein-coding sequence and its UTR is 6 times longer than the coding region. Furthermore, this UTR is not present in the transcript of any other species, and it is absent from the CFLAR gene of other primates. However, a sequence of 12kb length is unlikely to be absent from closely related species, thus we expanded our investigation to the flanking regions of the CFLAR gene. Through this analysis we found that other primates have the UTR vaguely conserved in the flanking regions of the CFLAR gene (**Fig7B**, See SI Fig.S6). This evidence suggests that the 12kb UTR of CFLAR is conserved and transcribed only in humans, thus its regulation by mt-miRNA_Met_ is probably possible only in humans.

**Fig 7.**
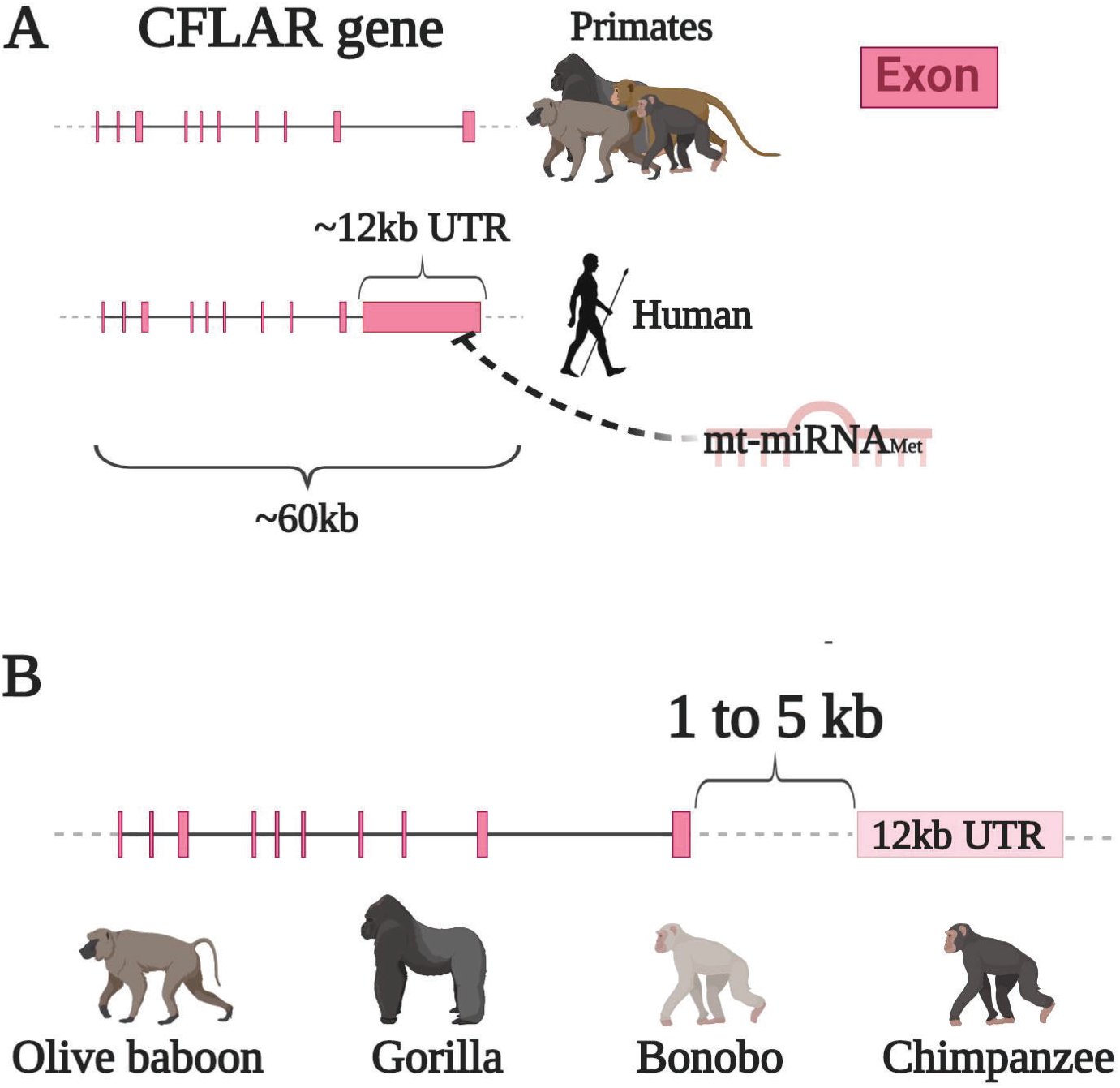
The mt-miRNA_Met_ can regulate CFLAR in humans only. In this figure, the genomic region of CFLAR is represented across multiple species, showing its conservation across primates **A**) A schematic representation of the genomic region of the CFLAR across multiple primates. We highlight the presence of this ~12kb UTR only in humans and show that the region binding mt-miRNA_Met_ is at the end of the UTR. **B**) We made a schematic representation showing that the ~12kb UTR present in humans is not translated anymore in primates but can be found at a short distance from the CFLAR gene (between 1 and 5 kb). No other miRNAs bind the CFLAR region bound by mt-miRNA_Met_. To see more details about the alignment of the genomic region of CFLAR between humans and other primates see Figure S6.

## Discussion

### The mtDNA is involved in RNA interference

Our study confirmed the presence of two shared features of nuclear miRNAs and mt-miRNAs, suggesting a role for the mtDNA in RNAi. The first shared feature is the ability to bind to Ago2. We provided definitive evidence that the mt-miRNAs can bind Ago2, the key protein in gene regulation through RNAi. Although we found some noise in the Ago2-IP samples, the genes having clearly defined mt-miRNAs were strongly upregulated in the comparison between Ago2-IP and mock-IP, thus supporting a genuine binding of these RNAs to Ago2. We verified the consistent presence of mt-miRNAs binding to Ago2 across multiple independent studies, suggesting that technical differences in RNA preparation, or sequencing, did not significantly affect the mt-miRNAs. Furthermore, our analyses uncovered clear transcriptional signatures across multiple genes and samples, confirming similar patterns observed in previous studies that analysed the expression patterns of small mitochondrial RNAs not bound to Ago2 (Pozzi et al. 2017; Pozzi and Dowling 2019; Mercer et al. 2011). The second shared feature is the conservation of an mt-miRNA across multiple species. This feature is present in many miRNAs (Lee, Risom, and Strauss 2007; Ambros 2004), and has been used before for phylogenetic purposes (Lee, Risom, and Strauss 2007; Sempere et al. 2006; Kenny et al. 2015). However, ours is the first study to demonstrate that some mt-miRNAs are conserved across multiple diverged species within Chordata. Although the conservation of miRNAs usually relies on verifying the presence of the miRNA sequence in the genome of the target species, this method was not possible for the mt-miRNAs because their sequence lies cryptic within the sequence of other host genes – in the case of mt-miRNA_Met_, this small RNA lies within the first half of the mt-tRNA Met gene. Thus, the presence of this sequence across multiple species only proves that the host gene is conserved. Nonetheless, by using small RNA expression data, we showed that the mt-miRNA_Met_ is expressed across multiple tissues and species with a conserved transcriptional signature. Arguably, this level of evidence is more reliable than the benchmark generally used for the identification of miRNAs, because we not only demonstrated the presence of the sequences within the genome, but also showed a clear and conserved transcriptional signature. Indeed, the conservation of these RNAs, in both sequence and expression, across species suggests a function, and the only known function of Ago2 is inhibition of mRNA translation (Ha and Kim 2014; Cloonan 2015). However, definitive proof that the mt-miRNAs act in RNAi in the same way as the miRNAs is still warranted. Indeed, to verify if the effect is similar, it would be necessary to know the target of specific mt-miRNAs and demonstrate their downregulation in protein expression through western blot analysis (Taylor et al. 2013). However, as a preliminary investigation of this phenomena, we used a mix of computational and experimental data to identify *bone fide* targets of the mt-miRNA that are most likely to be functional.

### The mt-miRNAMet - the selfish miRNA?

By investigating whether the mt-miRNA_Met_ targets any mRNA in the human brain, we uncovered evidence that mt-miRNA might be involved in the protection of mitochondria from apoptosis. We identified strong evidence that CFLAR is a target of mt-miRNA_Met_. CFLAR is a pseudo-caspase with multiple alternative transcripts known for having anti-apoptotic effect (S. A. Sarkar et al. 2009; He and He 2013). However, the mt-miRNA_Met_ targets only a specific isoform of CFLAR, named CFLAR_L_, since this is the only transcript transcribing the 12kb UTR hosting the Ago2 binding site. CFLAR_L_ is different from the other transcripts (Yu, Jeffrey, and Shi 2009; Tsuchiya, Nakabayashi, and Nakano 2015), and it is one of the most efficient activators of procaspase-8, a key factor in apoptosis (Chang et al. 2003). The activation of procaspase-8 is a known trigger of apoptosis, a cellular mechanism in which the mitochondria are destroyed, along with the hosting cell (Chang et al. 2003). This process is fundamental for eukaryotic organisms, however, it means extinction for the genetic material within the cell, thus this process could be negatively selected at cellular level, even if it was under positive selection at the organismal level (increased fitness of the individual). Intriguingly, a mitochondrion expressing an RNA such as mt-miRNA_Met_, would likely have an advantage relative to other mitochondria within a cellular environment if it was associated with a decreased possibility its host cell would die (Rand 2001). This is supported by the evidence that the mitochondrial genome is under strong selection (Ferguson and von Borstel 1992; MacAlpine, Perlman, and Butow 2000). Therefore, we hypothesize that this mt-miRNA might be protecting mitochondrial fitness at cellular level, thus being the first “selfish miRNA”, a miRNA expressed from an organelle which “protects” its own genome from a cellular mechanism.

### The mt-miRNAMet is involved in a human-specific regulation pathway

Our results suggest that the regulation of CFLAR by mt-miRNA_Met_ is probably unique to humans. Indeed, mt-miRNA_Met_ binds a UTR not transcribed in any other species. UTRs are not translated into proteins, and their role is usually to harbour regulatory sequences, thus, the presence of such a long UTR is puzzling (Yoon et al. 2008). Its presence, a long UTR sequence might be explained if this sequence harbours important regulatory sequences for this gene, such as the binding region for mt-miRNAMet. Our study supports this hypothesis, as a shortening or loss of this UTR, as seen in the other primates, would remove the binding site necessary for mt-miRNA_Met_ to regulate CFLAR expression, thus disrupting transcript regulation. Indeed, losing this regulatory region might cause an increase of CFLAR_L_ and potentially cell death through apoptosis (Tsuchiya, Nakabayashi, and Nakano 2015). Furthermore, our finding of a human-specific regulatory pathway for CFLAR aligns with the conflicting results found in experiments on CFLAR function (Tsuchiya, Nakabayashi, and Nakano 2015). In fact, most experiments have been performed in humans and mice, which although express the same protein, do not share the same regulatory region (Irmler et al. 1997; Shu, Halpin, and Goeddel 1997; Micheau et al. 2002). Indeed, mouse, like all other species, do not possess the 12kb UTR. Therefore, experiments on CFLAR function have conflicting results across humans and mice most likely due to the presence of species-specific regulation, such mt-miRNA_Met_ as in humans, supporting the finding that mt-miRNA_Met_ is part of a human-specific pathway.

### Are the mt-miRNAs acting outside the mitochondria?

Notwithstanding the ability of the mt-miRNAs to bind Ago2, their function is expected to be exerted outside the mitochondria. However, the mechanism that would lead the mt-miRNAs out of the mitochondria and into the cytoplasm remains unknown. The ability to bind Ago2 suggests the ability of mt-miRNAs to translocate outside to reach the protein, most likely using the same, yet not fully characterized, transport mechanism used by other nucleic acids. Indeed, proteins like the PNPase have been demonstrated to be involved in RNA translocation through the mitochondrial membrane (Wang et al. 2012). Furthermore, a recent study showed that small mtDNA fragments travel outside of the mitochondria through mitochondrial pores that might be used by small RNAs too, since they have very similar characteristics (Moriyama, Koshiba, and Ichinohe 2019). Our study is not the first supporting the ability of mt-miRNAs to move outside the mitochondria. Indeed, Maniataki and Mourelatos (2005) previously reported that the mitochondrial mRNA mt-tRNA Met (that we have shown here encodes mt-miRNA_Met_), is able to bind Ago2 outside the mitochondria. Furthermore, two other studies have previously provided evidence that Ago2 might be able to move inside of the mitochondria or to co-localize with it (Bandiera et al. 2011; Zhang et al. 2014). These studies performed western blots targeting Ago2 in samples with mitochondria-isolates, finding a positive signal for this protein. However, we believe the results of these studies better reflect co-localization of Ago2 and mitochondria than transport of Ago2 into the mitochondria. In fact, in the analyses of both Bandera et al. (2011) and Zhang et al. (2014), the Ago2 band was incredibly faint in the mitochondrial fractions, while other co-localization studies have revealed that Ago2 is very often co-localized with mitochondria (Olivieri et al. 2010; Vagin et al. 2013; Rogers et al. 2017). In sum, this indicates that Ago2 is being localized on the surface of the mitochondria, similarly to other proteins involved in RNAi (Vagin et al. 2013; Rogers et al. 2017). In this case, the evidence of mt-miRNA binding Ago2 suggests the presence of a mechanism enabling small RNAs transport through the mitochondrial membrane, potentially through PNPase (Dhir et al. 2018; D. Sarkar and Fisher 2006). Furthermore, we are aware of many types of small RNAs moving between mitochondria and cytoplasm, such as MitomiRs (Duarte, Palmeira, and Rolo 2015), nuclear miRNAs localized within the mitochondria; and double-stranded RNAs, acting in the cytoplasm to trigger an immune response (Dhir et al. 2018). The mt-miRNAs might use the same, as-yet uncharacterized transport mechanism used by these other RNAs.

### A new class of RNA without a new name

Small mitochondrial RNAs have been previously reported and given multiple new names across a series of studies. The first study reporting their existence did not explicitly name these RNAs, simply mentioning the presence of highly expressed small mitochondrial RNAs (Mercer et al. 2011). In 2013, Ro *et al.* showed that small mitochondrial RNAs were encoded in both mouse and humans, potentially having a function in mitochondrial regulation (Ro et al. 2013). This study named these RNAs as ‘mitosRNAs’, however, they annotated thousands of RNAs in this new class, in which the described RNAs had very different sizes (from ~15nt to ~120nt), were barely expressed, and had no associated evidence of function. In 2017, another group tried to name these RNAs using other standards (based on small RNAs of high expression, they called them ***sm***all ***h***ighly ***t***ranscribed small RNAs; smithRNAs), but introducing further confusion as to the nomenclature of these RNAs (Pozzi et al. 2017). This confusion persists due to the lack of precise standards when classifying new functional RNAs. Therefore, we have decided to keep the nomenclature simple and reasonable, adhering to the historical precedent, and following the example set by the authors discovering one of the most famous classes of small non-coding RNAs: the piRNAs (Grivna et al. 2006; Kim 2006). The discoverer of the piRNAs used their ability to bind the protein Piwi as the criterion to define them. Thus, we classify only those small mitochondrial RNAs able to bind Ago2 as mt-miRNAs. Furthermore, as mentioned above, we believe that further increasing the RNA nomenclature would not benefit the scientific community (Ro et al. 2013; Pozzi et al. 2017; Srinivasan and Das 2015), hence we propose adding a simple prefix (mt-) to define this class of small RNAs (Budak et al. 2016). Along with tRNAs and rRNAs, which receive the prefix mt-when referring to those encoded by mtDNA, we contend that miRNAs that bind Ago2 should similarly receive the same prefix when encoded in the mtDNA.

### Some mt-miRNAs are inexplicably long

Some mt-miRNAs are almost twice the length of nuclear miRNAs but still able to bind Ago2, a case never seen before. Given that the protein complex necessary for the RNAi usually binds RNAs that are 20-30nt long (Ha and Kim 2014; Cloonan 2015), it remains unclear if longer mt-miRNAs will have similar functions once bound to Ago2. Some small RNAs in *C. elegans* are 34nt long, longer than most mt-miRNAs (~32nt), and are able to bind another protein involved in RNAi, Piwi (Ha and Kim 2014; Cloonan 2015)4). However, no previous miRNAs have been reported as long as the long isoform of mt-miRNA Thr (41nt), which usually would suggest this mt-miRNA is an intermediate precursor of the mt-miRNA. However, our analysis in paired long and short mt-RNAs found no support for this precursor hypothesis, due to the lack of longer RNAs in the mt-tRNA Thr. We suggest this RNA might be associated with a different function from canonical miRNAs. Indeed, in studies using Ago2-IP samples, the RNAs of ~35 length are usually considered fragments of miRNA targets, such as mRNAs, and not miRNAs (Boudreau et al. 2014). The hypothesis that the mt-tRNA Thr might be a target of Ago2 is supported by our analysis of mt-miRNAs in RNA-seq samples without Ago2-IP treatment in mouse. In these samples, we found clearly defined transcriptional signatures in several mt-tRNAs, such as mt-tRNA Ser 1, but not in the mt-tRNA Thr, suggesting that the mt-tRNA Thr does not encode for any small RNA and what was found in the Ago2-IP samples is a fragment of a longer RNA bound to Ago2. However, the samples analysed in both studies come from different tissues, thus it is possible that lack of mt-miRNAs from mt-tRNA Thr results from tissue-specific expression. The presence of long mt-miRNAs or the binding of Ago2 to these mt-tRNAs is puzzling, and without any obvious explanation. Nonetheless, this result shows that the mt-miRNAs and the miRNAs do not share a similar biogenesis, as the mt-miRNAs are not matured from ~70nt pre-miRNAs as are the nuclear miRNAs.

### Multiple gene layers in the mitochondria

Our study shows that the mtDNA harbours multiple gene layers, and that the products of overlapping genes are selected during the primary mt-RNA maturation. We demonstrated that the mt-miRNAs are encoded within other genes, specifically protein-coding genes or mt-tRNAs, aligning to observations of previous studies (Pozzi et al. 2017; Pozzi and Dowling 2019; Ro et al. 2013; Riggs et al. 2018; Bottje et al. 2017; Mercer et al. 2011; Larriba, Rial, and Del Mazo 2018). Likewise, other mitochondrial products are encoded within multiple genes: double-stranded RNAs (Dhir et al. 2018), long non-coding RNAs (Rackham et al. 2011), and proteins (C. Lee et al. 2015; C. Lee, Yen, and Cohen 2013; K. H. Kim et al. 2018). However, some of these mitochondrial products have been better described than others. The double-stranded RNAs have been discovered only very recently, and function in the immune response (Dhir et al. 2018). The function of long non-coding RNAs is still unknown, but their nuclear counterparts are known for having extensive roles in gene expression regulation (Mattick 2003). There are also several newly discovered mitochondrial proteins, with ‘humanin’ the first and most intensely studied (Hashimoto et al. 2001). This protein is encoded within rRNA 16S and, although its function is not fully understood, it seems to somehow have protective effects against Alzheimer’s disease (Matsuoka 2009). Thus, the number of functional mitochondrial products identified in recent years is quickly increasing, providing strong support for the presence of multiple new regulatory layers within the mitochondrial genome. These findings are intriguing because the presence of these novel mitochondrial products suggests a reinterpretation of the candidate mechanisms by which pathogenic mutations in the mtDNA sequence exert their effects on organismal health and function may be required in several cases.

### The mt-miRNAs change our perspective on mitonuclear interactions

The mt-miRNAs add another level of complexity to the dynamics of mitonuclear communications. Indeed, because the mt-miRNAs share features with the miRNAs, we expect them to affect cell biology in a similar manner. The miRNAs play pervasive roles in cell regulation (Friedman et al. 2009), and the presence of mt-miRNAs indicates that the mtDNA might broadly affect cell regulation as well. The mt-miRNAs might act as a vector to affect nuclear regulation in many ways. Indeed, by using sequence complementarity, mt-miRNAs could lead Ago2 to interfere with the gene expression of virtually any nuclear mRNA that exhibits partial sequence complementarity (Cloonan 2015; Ambros 2004). This could help explain many of the diverse phenotypes linked to mtDNA mutations observed in recent years (Hopkins et al. 2017; Hudson et al. 2014; Dobler et al. 2014). Recent studies, for example, have demonstrated clear associations of mtDNA mutations on a range of phenotypes, ranging from thermal tolerance, to cognitive function, to fertility (Lajbner et al. 2018; Camus et al. 2017; Yee, Sutton, and Dowling 2013; Dowling, Abiega, and Arnqvist 2007; Roubertoux et al. 2003), as well as a range of human diseases not previously associated with mitochondrial genetics (Hopkins et al. 2017; Hudson et al. 2014). However, the mechanisms underpinning these diverse effects associated with the mitochondrial genome are yet to be understood. We believe that RNAi, mediated through mt-miRNAs, might well provide the explanation for the diversity of phenotypic effects associated with mitochondrial sequence variation.

### The unusual maturation of the mt-miRNAs

The mitochondria generally transcribe long mt-RNA precursors encoding multiple genes (D’Souza and Minczuk 2018), potentially including the mt-miRNAs. Indeed, the transcriptional signature of the mt-miRNAs does not match the transcriptional signature of the long mitochondrial RNAs (~70nt), suggesting that the mt-miRNAs are matured directly from the pre-mt-RNAs. Mitochondrial genes are usually transcribed in few long polycistronic mt-RNA, RNAs including multiple genes, and these RNAs are called pre-mt-RNAs (Van Haute et al. 2015). Although previously the presence of intermediate RNAs were proposed for the mt-miRNAs (Pozzi et al. 2017), our results suggest that the mt-miRNAs originate from the pre-mt-RNAs. Indeed, we identified multiple cases of specific isoforms of mt-miRNAs whose sequence spanned two genes. In these cases, it is clear that miRNAs cannot originate from a mature tRNA, since they would then lack part of the sequence. Furthermore, mt-tRNAs undergo a vast number of nucleotide modifications through their maturation (Richter et al. 2018; Pan 2018), thus mt-miRNAs originating from mature tRNAs will have these RNA modifications, thus escaping normal sequencing methods. That is, it would be impossible for the mt-miRNAs presented in our analyses to exhibit these modified nucleotides, given the methods used for the RNA sequencing would not have captured these modified RNAs (Zheng et al. 2015). Therefore, according to our results, the mt-miRNAs encoded in mt-tRNAs are either matured from them before the posttranscriptional modifications, or directly from the mt-RNA precursor. Our study provides the first clues as to the biogenesis of the mt-miRNAs and, in general, supports the hypothesis that the mitochondria regulates its products mostly from the pre-mt-RNA (Sloan and Wu 2016; Lavrov et al. 2016).

### Overlapping selection

Mitochondrial genes are known to be under constant strong purifying selection (J. W. Ballard and Kreitman 1995; Ballard and Whitlock 2004), however, the existence of multiple gene layers suggests the presence of overlapping selection pressures that could alter the strength or direction of selection on particular regions of mtDNA sequence. Indeed, theory suggests that multiple products encoded within the same region of a gene would affect the selection pressure on this region (Rogozin et al. 2002). We hypothesise that the presence of overlapping genes such as mt-miRNAs and dsRNAs (Dhir et al. 2018), on canonical genes such as mt-tRNAs, will increase the effect of purifying selection in order to preserve the function of these products. Our results support this hypothesis, showing that within the mt-tRNA Met, the region harbouring the mt-miRNAMet has fewer polymorphisms than did the rest of the tRNA gene. While more analysis across a broad number of species and mt-miRNAs will be necessary to fully test this hypothesis, this study provides the first evidence that this phenomenon might exist.

## Author Contributions

A.P. and D.K.D. conceived the study. A.P. performed the analyses. A.P. and D.K.D. discussed the results and wrote the manuscript.

## Acknowledgement

We wish to acknowledge all researchers who made their data freely available, thus giving us the chance to perform this work. The study was funded by the Australian Research Council, through grants DP170100165 and FT160100022 to D.K.D, and through Australian Government Research Training Program (RTP) Scholarship to A.P.

## CONTACT FOR REAGENT AND RESOURCE SHARING

Further information and request about the reagents can be directed to the Lead Contact, Andrea Pozzi. However, as the data are sourced from previously published articles, we can only redirect you to the appropriate source, and we cannot personally provide full details of each reagent.

## EXPERIMENTAL MODEL AND SUBJECT DETAILS

This study is based on published data deposited in the Sequence Read Archive (SRA) in NCBI, thus all information listed here are sourced from the original study where the data were obtained.

### Cell lines

According to the original authors (Zhang et al., 2018), HeLa cells were grown on 15 cm plates using MEDM plus 10% FBS. The neuronal progenitor (NP) and teratoma-derived fibroblast (TDF) cell culture, were derived from human embryonic stem cells (hESC). The differentiation of hESC into NP was induced by replacing the original growing medium with DMEM/F12 supplemented with 2% B27, 100ng/ml FGF, 100ng/ml EGF and 5ng/ml heparin. Further details on the methods can be found at GSE115146 in the Gene Expression Omnibus (GEO) database and in the original article.

The differentiation of hESC into TDF was induced by injection of resuspended cells into mice homozygous for severe combined immune deficiency spontaneous mutation (SCID). The tumors grew in after 6 weeks were then removed and cultured in a medium of 10% FBS, nonessential amino acids, 2mM glutamine, 1% penicillin/streptomycin and 0.55μM β-mercaptoethanol. Further details on the methods, as described by the original authors, can be found at GSE112006 in the Gene Expression Omnibus (GEO) database.

According to the original authors (Zamudio et al., 2014), the embryonic stem cell culture (mESC) were grown on gelatinized tissue culture plates in Dulbecco’s Modified Essential Media supplemented with multiple other nutrients. The full list of nutrients and details about the growing method can be found at GSE50595 in the GEO database and in the original article.

### Model organisms

According to the original authors (Jee et al., 2018; Viljetic et al., 2017), the experiments on mice sourced from the Jackson laboratory were carried out in compliance with their institutional protocols, thus we believe that correct ethics for the experiments have been followed.

According to the original authors (Woo et al., 2017), the experiments on mice sourced by Stefan Somlo were carried out in compliance with the Animal Care and Use Committee (IACUC) rules at Sookmyung Women’s University. Therefore, we believe that correct ethics were followed during the experiment. According to the original authors, the samples from the chickens were sourced by Peter Jensen, however, there is not any further details about how the chickens were housed and maintained.

According to the original authors (Vaz et al., 2015), the zebrafish (Singapore strain) were maintained according to the Animal Care and Use Committee (IACUC) rules. Therefore, we believe that correct ethics were followed during the experiment.

### Human samples

The BA9 samples were obtained by the original authors (Hoss et al., 2015), and the gathering of the samples were exempt from ethics approval because the study involves only tissue collected post-mortem, and consequently not classified as human subjects. This decision was taken by the Boston University School of Medicine Institutional Review Board (Protocol H-28974).

The CHC samples were obtained by the original authors (Butt et al., 2016), and the original authors declared that written informed consent for the use of biological samples and clinical records was given by all the participants. Furthermore, they declared that their work was done in accordance with the ethical guidelines of the 1975 Declaration of Helsinki and the International Conference on Harmonization Guidelines for Good Clinical Practice.

The TL samples were obtained by the original authors (Maragkakis et al., 2016; Nakaya et al., 2013), and the authors provided only a brief description of the samples. Indeed, they only mention that the samples come from temporal surgical lobectomy of three unrelated individuals. Although no statements on ethics is present in the original articles, we believe that is likely that the correct ethics approval processes are followed in the institution where the experiments where performed (University of Pennsylvania).

## METHOD DETAILS

### Ago2 co-immunoprecipitation sequencing

According to the original authors (Zhang et al., 2018), before the Ago2 immunoprecipitation (IP) of HeLa cells, the cultures were UV irradiated at 400mj. The IP was performed using Anti-Ago2 (Abnova, H00027161-M01), and following the IP the RNA libraries were made and sequenced using Hi-Seq 2500. Further details can be found in the original article and in the GEO database (GSE115146).

According to the original authors, before the Ago2-IP of the NP and TDF, the cultures were UV irradiated at 400mj. The IP was performed using Anti-Ago2 (ProteinTech) and following the IP the RNA libraries were made and sequenced using Hi-Seq 2000. Further details can be found in the GEO database (GSE112006).

According to the original authors (Zamudio et al., 2014), the mESC cells have been modified to leave only a modified Ago2 active in these cells. This modified Ago2 gene is known as FLAG-hemagglutinin (HA)-tagged hAgo2 (FHAgo2), thus has a specific epitope that can be targeted for IP. Furthermore, contrary to the experiments in other cell lines, in this case the cell cultures were lysed before the UV-cross linking and IP. Further details can be found in the original article and in the GEO database (GSE50595).

The Ago2-IP of the TL samples were performed using 2A8 Anti-Ago monoclonal antibody tethered to Protein A Dynabeads (Invitrogen), according to the original authors (Maragkakis et al., 2016). Further details can be found in the original article and in the BioProject NCBI database (PRJNA299324).

### Ago2 immunoprecipitation miRNA targets enrichment

The original authors of the study performed the mRNA target enrichment (Maragkakis et al., 2016), and we describe here a short summary of their method. After the Ago2-IP of the three brain samples, the authors isolated longer RNAs (miRNA targets) using nitrocellulose filter, and 8% PAGE gels. Once the targets were purified, they have been re-amplified by PCR and sequenced. Further information about the methods can be found in previous works of the authors (Maragkakis et al., 2016; Nakaya et al., 2013).

### Mt-miRNAs sequence alignment

The alignment of the RNA-seq libraries, both with and without IP, were performed using BowTie2 (Langmead & Salzberg, 2013). We did not remove the adapter for each library, and instead used the soft clipping provided by bowtie2 through the setting *--local*. Due to the possible differences between the reference genome used, and the sequence of the individual used for analysis, we used non-stringent parameters in the alignment. Thus, for the alignment we set a seed of 20 nucleotides (*−L 20*) and we allowed up to 1 mismatch (*−N 1*) between the small RNA and the reference genome. To gauge the overall amount of small RNAs within one gene, we used samtools *idxstats* function (Li et al., 2009), which outputs the reads aligning to each chromosome. Likewise, to obtain the coverage for each nucleotide, and thus identify the mitochondrial genes having the transcriptional signature of a mt-miRNA, we used the function *genomecov* of bedtools (Quinlan & Hall, 2010). To generate the total coverage for each gene, we used the setting *genomecov -d -i* while for the 5’ and 3’ coverage we added the setting *−5* and *−3* respectively. To generate the correct format to plot the gene coverage in the R package *circlize*, we used the parameter *-bg* instead of *-d* on bedtools *genomecov*. The alignment was done using the current reference genomes for human (NC_012920.1) and mouse (NC_005089.1).

### mt-miRNA_Met_ polymorphism identification

The identification of polymorphism on the mt-miRNA_Met_ across multiple species was performed by visual comparison of the sequences downloaded from the mt-tRNAs database mitotRNAdb (http://mttrna.bioinf.uni-leipzig.de/mtDataOutput/Welcome). We only used the reference sequences to establish the presence of polymorphisms, and we did not include any population data, because the quality of population data (i.e. the frequency of specific polymorphisms) are very different across the organisms considered. Therefore, we used the reference genome present in mitotRNAdb for *Danio rerio* (NC_002333.2), *Gallus gallus* (NC_001323.1), *Homo sapiens* (NC_012920.1), and *Mus musculus* (NC_005089.1).

### mt-miRNA_Met_ target prediction

The *in silico* prediction of the mt-miRNA_Met_ target was done using the web-server MR-microT (Kanellos et al., 2014; Reczko et al., 2012), a program able to use a custom miRNA sequence to identify potential target mRNAs in the human genome (using the database Ensembl v84). This program outputs both a score, indicating how likely is the predicted target to be real, and the regions of the sequence where miRNA and mRNAs are predicted to interact. As mentioned in the results, the list of potential targets predicted by this program is 8,709, which we filtered using the gene ontology (GO) term Insulin Response (GO:0032869) to obtain a list of 74 potential targets. To experimentally verify these targets, we extracted the coverage of each gene using bedtools (Quinlan & Hall, 2010) with settings *genomecov -d -i*, and then inspected by eye for the presence of sequences in the regions corresponding to the prediction of MR-micro T. Although this method might be missing several targets of this gene (false negative), we focused on finding targets that have the highest chance of being real (true positive). It is worth noting that the mRNA target identified using this method, CFLAR, has a MR-micro T score with high likelihood to be predictive (>0.8) according to the authors of the program (Kanellos et al., 2014; Reczko et al., 2012).

### Comparative analysis mt-miRNA_Met_ and nuclear miRNAs

The miRNAs were quantified using the output of samtools *flagstat* option, which provides the number of aligned reads, both the total amount and as a percentage of the RNA pool, for the specified reference sequence. We used mt-tRNA Met as the reference sequence to quantify the amount of mt-miRNA Met reads expressed, indeed, over 90% of the reads come from the region of the mt-miRNA_Met_, thus using the coverage of the entire mt-tRNA Met provides a very good approximation of mt-miRNA_Met_ without excluding any of its isoforms. To identify miRNAs that might be binding the same regions as mt-miRNA_Met_, we used TargetscanHuman 7.2 (http://www.targetscan.org/) using CFLAR as a query, and then we annotated all the miRNAs predicted to bind the same, or similar, positions of mt-miRNA_Met_. Using this approach, we annotated 13 miRNAs: hsa-miR-6747-3p, hsa-miR-6778-3p, hsa-miR-150-5p, hsa-miR-186-3p, hsa-miR-4698, hsa-miR-4731-5p, hsa-miR-5589-5p, hsa-miR-6506-5p, hsa-miR-619-5p, hsa-miR-5089-5p, hsa-miR-6504-3p, hsa-miR-4438, hsa-miR-5095, hsa-miR-7151-3p. To quantify the nuclear miRNAs, we used the sequences present in miRBase (http://www.mirbase.org/) as a reference sequence for *samtools*. However, none of these nuclear miRNAs was found in brain tissues. To verify the absence of artifacts, or errors in our analysis, we used the same method to quantify common, well-known miRNAs. The miRNAs we chose are hsa-Let7a-1, hsa-miR-9-5p, hsa-miR-100. The similar abundance of these miRNAs to the mt-miRNA_Met_ confirmed the likely absence of artifacts or errors in the analysis, thus confirming the absence of other known miRNAs binding the same position of mt-miRNA_Met_.

### CFLAR Untranslated region analysis

We verified the presence of the 3’ untranslated region (UTR) of 20kb length found in *Homo sapiens* across multiple species by searching on the online version of blastn using default settings (https://blast.ncbi.nlm.nih.gov/Blast.cgi?PAGE_TYPE=BlastSearch). After not finding any hit on blastn, we tested the presence of this UTR across several species in the Gene database of NCBI (https://www.ncbi.nlm.nih.gov/gene). Due to the lack of this UTR across any other species, we focused more detailed analysis on species closer to *Homo sapiens* as they are more likely to have some trace of this sequence. Thus, we selected five other high-quality genomes from different primates and compared the full genomic region of CFLAR. The species involved are *Homo sapiens* (ID: 8837), *Pan troglodytes* (ID: 459872), *Pan paniscus* (ID: 100972044), *Pongo abelii* (ID: 100172025), *Gorilla gorilla* (ID: 101130329), and *Papio anubis* (ID: 101006231). The alignment of the different regions was made using LastZ (http://www.bx.psu.edu/~rsharris/lastz/), with settings *--step=10 --nogapped --ambiguous=iupac --matchcount=20 --format=rdotplot*, which provided a matrix of the alignment that was then used for the dotplot in R.

