## Supplemental Figure 1 for "A new member in the Argonaute crew: the mt-miRNAs"

### Gene-by-gene analysis of samples treated with different contrations of RNase I

The **figure S1** highlights the lack of effect from different concentrations of RNase I, across multiple mitochondrial genes.

In the following figure, we show the transcriptional signatures of putative small RNAs across all mitochondrial genes, gene by gene. The gene order is protein-coding, rRNAs, tRNAs, and within each group, the genes are in alphabetic order. In the following paragraph, we highlight the genes encoding mt-miRNAs, using the same prerequisite criteria as in the manuscript (i.e. transcriptional signatures with clear cut ends across multiple biological replicates indicate the presence of an mt-miRNA). Among the protein-coding genes, none of the genes has mt-miRNAs fitting the rationale for mt-miRNAs identification. Among the genes encoding for the two mt-rRNAs, only the mt-rRNA12s have an mt-miRNAs encoded at position 250 which is consistently transcribed across multiple biological replicates and have well-defined start and end. Among the mt-tRNA genes, only mt-tRNA Phe (trnF), mt-tRNA Met (trnM), mt-tRNA Gln (Q), and mt-tRNA Trp (trnW) are consistently transcribed across multiple biological replicates and have well-defined start and end.

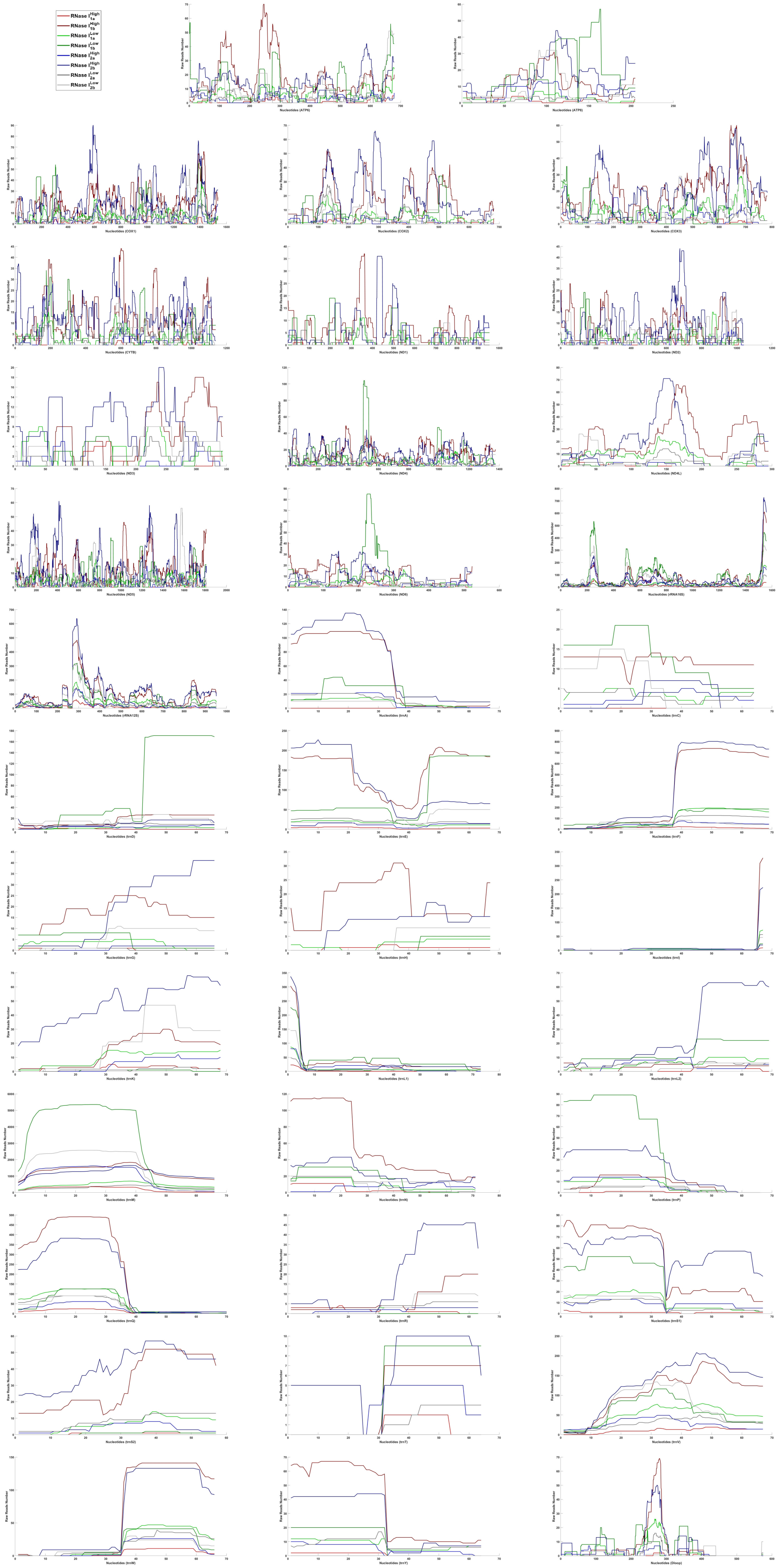

RNase I High 1a
RNase I High 1b
RNase I Low 1a
RNase I Low 1b
RNase I High 2a
RNase I High 2b
RNase I Low 2a
RNase I Low 2b
Raw Reads Number
Nucleotides (ATP6)
Nucleotides (ATP8)
Nucleotides (COX1)
Nucleotides (COX2)
Nucleotides (COX3)
Nucleotides (CYTB)
Nucleotides (ND1)
Nucleotides (ND2)
Nucleotides (ND3)
Nucleotides (ND4)
Nucleotides (ND4L)
Nucleotides (ND5)
Nucleotides (ND6)
Nucleotides (rRNA16S)
Nucleotides (rRNA12S)
Nucleotides (trnA)
Nucleotides (trnC)
Nucleotides (trnD)
Nucleotides (trnE)
Nucleotides (trnF)
Nucleotides (trnG)
Nucleotides (trnH)
Nucleotides (trnI)
Nucleotides (trnK)
Nucleotides (trnL1)
Nucleotides (trnL2)
Nucleotides (trnM)
Nucleotides (trnN)
Nucleotides (trnP)
Nucleotides (trnQ)
Nucleotides (trnR)
Nucleotides (trnS1)
Nucleotides (trnS2)
Nucleotides (trnT)
Nucleotides (trnV)
Nucleotides (trnW)
Nucleotides (trnY)
Nucleotides (Dloop)
