## Supplemental Figure 2 for "A new member in the Argonaute crew: the mt-miRNAs"

### Gene-by-gene analysis of Ago2-IP and mock-IP samples

Fi**gure S2** highlights the enrichment of Ago2-IP samples compared to IgG-IP samples, across multiple mitochondrial genes.

In the following figure, we show the transcriptional signature of small RNAs across all mitochondrial genes, divided by gene for neuronal progenitors (NP) and teratoma-derived fibroblast (TDF). The gene order is protein-coding, rRNAs, tRNAs, and within each group, the genes are in alphabetic order. In the following paragraph, we highlight the genes encoding for mt-miRNAs, following the same prerequisites as outlined in the manuscript (that transcriptional signatures with clear cut ends across multiple biological replicates indicate the presence of an mt-miRNA).

***In neuronal progenitors (NP)***

Among the protein-coding genes, none of the genes has mt-miRNAs fitting the prerequisites for mt-miRNA identification. Among the genes encoding for the two mt-rRNAs, none of the genes has an mt-miRNA consistently transcribed across multiple biological replicates and a well-defined start and end. Among the mt-tRNA genes, only mt-tRNA Phe (trnF), mt-tRNA Met (trnM), mt-tRNA Ser 2 (trnS2), and mt-tRNA Tyr (trnY) have mt-miRNAs consistently transcribed across multiple biological replicates and having well-defined start and end.

***In teratoma-derived fibroblast (TDF)***

Among the protein-coding genes, only ATP8 encodes for an mt-miRNAs fitting the rationale for mt-miRNAs identification. Among the genes encoding for the two mt-rRNAs, none of the genes has an mt-miRNA consistently transcribed across multiple biological replicates and has a well-defined start and end. The peaks are due to the overlap of many different small RNAs partially overlapping. Among the mt-tRNA genes, only mt-tRNA Glu (trnE), mt-tRNA Phe (trnF), mt-tRNA Met (trnM), mt-tRNA Gln (trQ), mt-tRNA Ser 1 (trnS1), mt-tRNA Ser 2 (trnS2), mt-tRNA Trp (trnW) and mt-tRNA Tyr (trnY) have mt-miRNAs consistently transcribed across multiple biological replicates and having well-defined start and end.

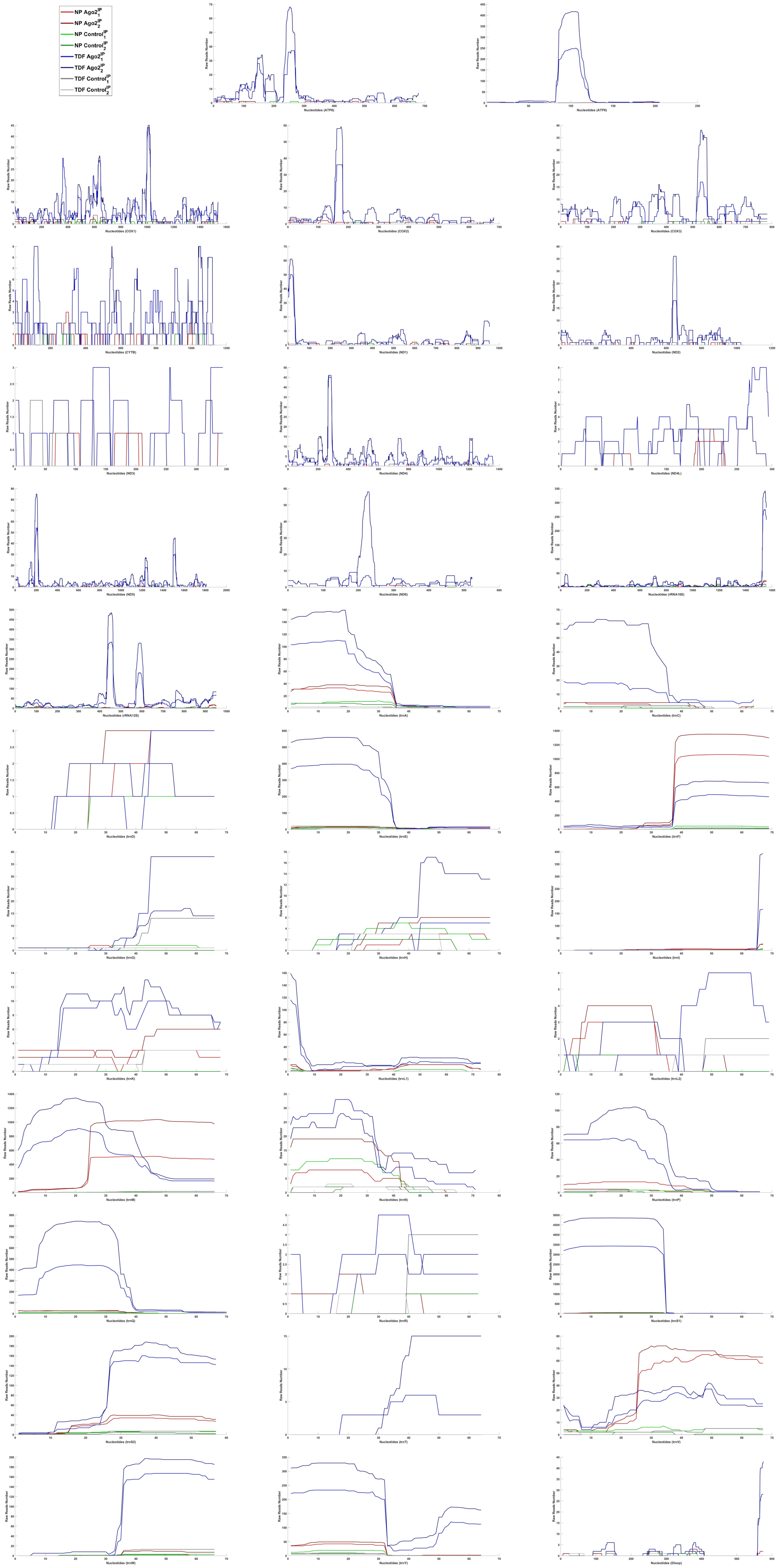
NP Ago2$_1^{IP}$
NP Ago2$_2^{IP}$
NP Control$_1^{IP}$
NP Control$_2^{IP}$
TDF Ago2$_1^{IP}$
TDF Ago2$_2^{IP}$
TDF Control$_1^{IP}$
TDF Control$_2^{IP}$
Raw Reads Number
Nucleotides (ATP6)
Nucleotides (ATP8)
Nucleotides (COX1)
Nucleotides (COX2)
Nucleotides (COX3)
Nucleotides (CYTB)
Nucleotides (ND1)
Nucleotides (ND2)
Nucleotides (ND3)
Nucleotides (ND4)
Nucleotides (ND4L)
Nucleotides (ND5)
Nucleotides (ND6)
Nucleotides (rRNA16S)
Nucleotides (rRNA12S)
Nucleotides (trnA)
Nucleotides (trnC)
Nucleotides (trnD)
Nucleotides (trnE)
Nucleotides (trnF)
Nucleotides (trnG)
Nucleotides (trnH)
Nucleotides (trnI)
Nucleotides (trnK)
Nucleotides (trnL1)
Nucleotides (trnL2)
Nucleotides (trnM)
Nucleotides (trnN)
Nucleotides (trnP)
Nucleotides (trnQ)
Nucleotides (trnR)
Nucleotides (trnS1)
Nucleotides (trnS2)
Nucleotides (trnT)
Nucleotides (trnV)
Nucleotides (trnW)
Nucleotides (trnY)
Nucleotides (Dloop)
