## Supplemental Figure 3 for "A new member in the Argonaute crew: the mt-miRNAs"

### Gene-by-gene analysis of HeLa cells with Ago2-IP and without

**Figure S3** highlights the enrichment in mt-miRNAs of specific genes after Ago2-IP treatment, across multiple mitochondrial genes.

In the following figure, we show the small RNAs transcriptional signature of all the mitochondrial genes, divided by gene. The gene order is protein-coding, rRNAs, tRNAs, and within each group, the genes are in alphabetic order. In the following paragraph, we highlight the genes encoding for mt-miRNAs, following the prerequisite condition that transcriptional signatures with clear cut ends across multiple biological replicates indicate the presence of an mt-miRNA. The samples are paired, thus the samples of each treatment have to be compared to their corresponding number. For example, $\mathrm{Input}_1$ should be compared to $\mathrm{Ago2}^{\mathrm{IP}}_1$ and not to $\mathrm{Ago2}^{\mathrm{IP}}_2$.

Among the protein-coding genes, none of the genes has mt-miRNAs fitting our criteria for mt-miRNA identification. Among the genes encoding for the two mt-rRNAs, none of the genes has an mt-miRNA consistently transcribed across multiple biological replicates and a well-defined start and end. Among the mt-tRNA genes, only mt-tRNA Ile (trnI), mt-tRNA Glu (trnQ), and mt-tRNA Ser1 (trnS1) are consistently transcribed across multiple biological replicates and have well-defined start and end.

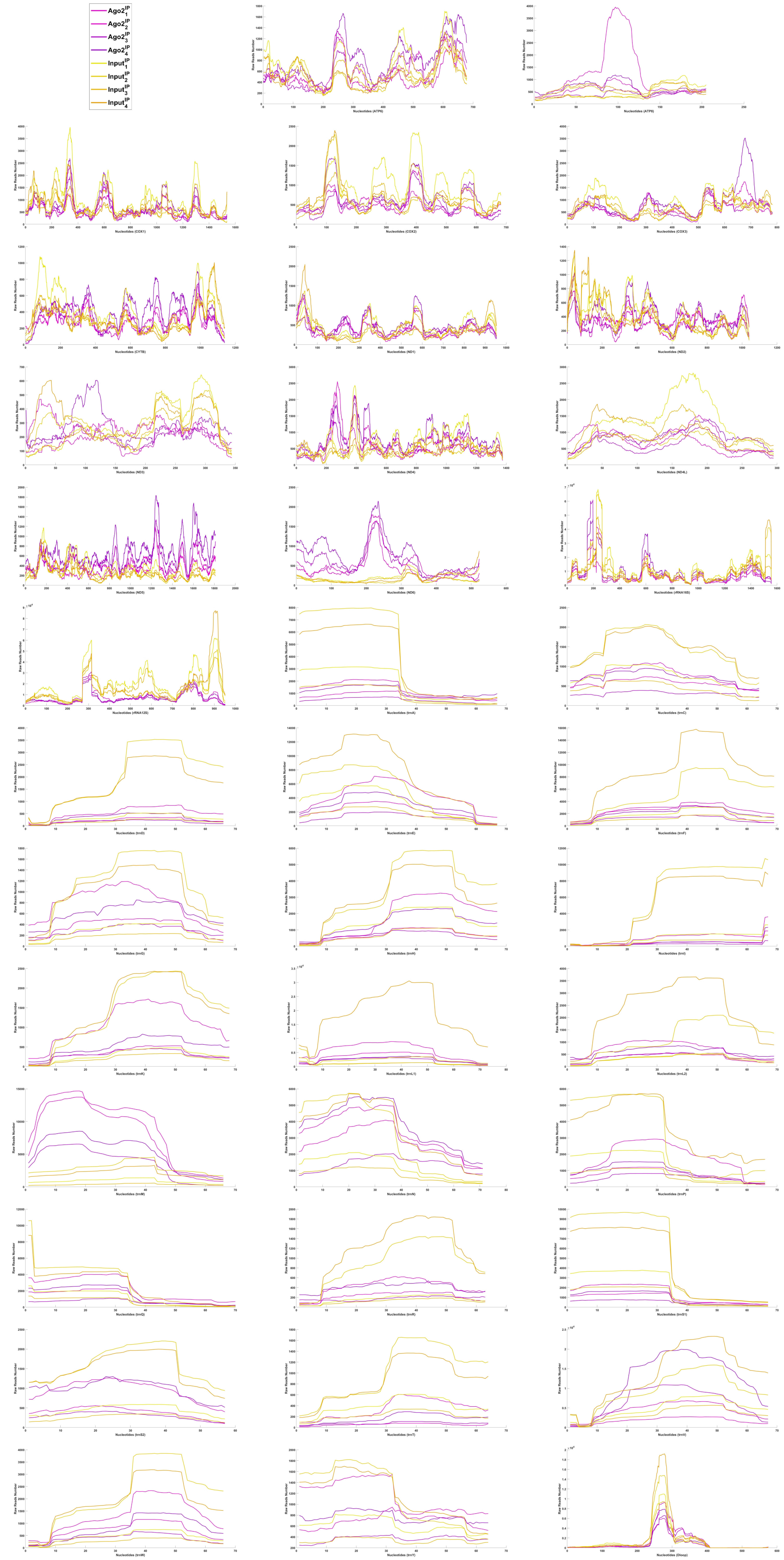
$Ago2_1^{IP}$
$Ago2_2^{IP}$
$Ago2_3^{IP}$
$Ago2_4^{IP}$
$Input_1^{IP}$
$Input_2^{IP}$
$Input_3^{IP}$
$Input_4^{IP}$
Raw Reads Number
Nucleotides (ATP6)
Nucleotides (ATP8)
Nucleotides (COX1)
Nucleotides (COX2)
Nucleotides (COX3)
Nucleotides (CYTB)
Nucleotides (ND1)
Nucleotides (ND2)
Nucleotides (ND3)
Nucleotides (ND4)
Nucleotides (ND4L)
Nucleotides (ND5)
Nucleotides (ND6)
Nucleotides (rRNA16S)
Nucleotides (rRNA12S)
Nucleotides (trnA)
Nucleotides (trnC)
Nucleotides (trnD)
Nucleotides (trnE)
Nucleotides (trnF)
Nucleotides (trnG)
Nucleotides (trnH)
Nucleotides (trnI)
Nucleotides (trnK)
Nucleotides (trnL1)
Nucleotides (trnL2)
Nucleotides (trnM)
Nucleotides (trnN)
Nucleotides (trnP)
Nucleotides (trnQ)
Nucleotides (trnR)
Nucleotides (trnS1)
Nucleotides (trnS2)
Nucleotides (trnT)
Nucleotides (trnV)
Nucleotides (trnW)
Nucleotides (trnY)
Nucleotides (Dloop)
