## Supplemental Figure 4 for "A new member in the Argonaute crew: the mt-miRNAs"

### Gene-by-gene analysis of mouse embryonic stem cells with Ago2-IP and without

**Figure S4** highlights the enrichment in mt-miRNAs of specific genes after Ago2-IP treatment, across multiple mitochondrial genes.

In the following figure, we show the small RNAs transcriptional signature of all the mitochondrial genes, divided by gene. The gene order is protein-coding, rRNAs, tRNAs, and within each group, the genes are in alphabetic order. In the following paragraph, we highlight the genes encoding for mt-miRNAs, following the prerequisite condition that transcriptional signatures with clear cut ends across multiple biological replicates indicate the presence of an mt-miRNA. The samples are paired, thus the samples of each treatment have to be compared to their corresponding number. For example, $InputIP_1$ should be compared to $Ago2^{IP}{}_1$ and not to $Ago2^{IP}{}_2$.

Among the protein-coding genes, only COX1, COX3, and ND4L have mt-miRNAs our criteria for mt-miRNA identification. Among the genes encoding for the two mt-rRNAs, none of the genes has an mt-miRNA consistently transcribed across multiple biological replicates and a well-defined start and end. Among the mt-tRNA genes, only mt-tRNA Glu (trnE), mt-tRNA Phe (trnF), mt-tRNA Leu2 (trnL2), mt-tRNA Met (trnM), and mt-tRNA Thr (trnT) have mt-miRNAs consistently transcribed across multiple biological replicates and showing well-defined start and end.

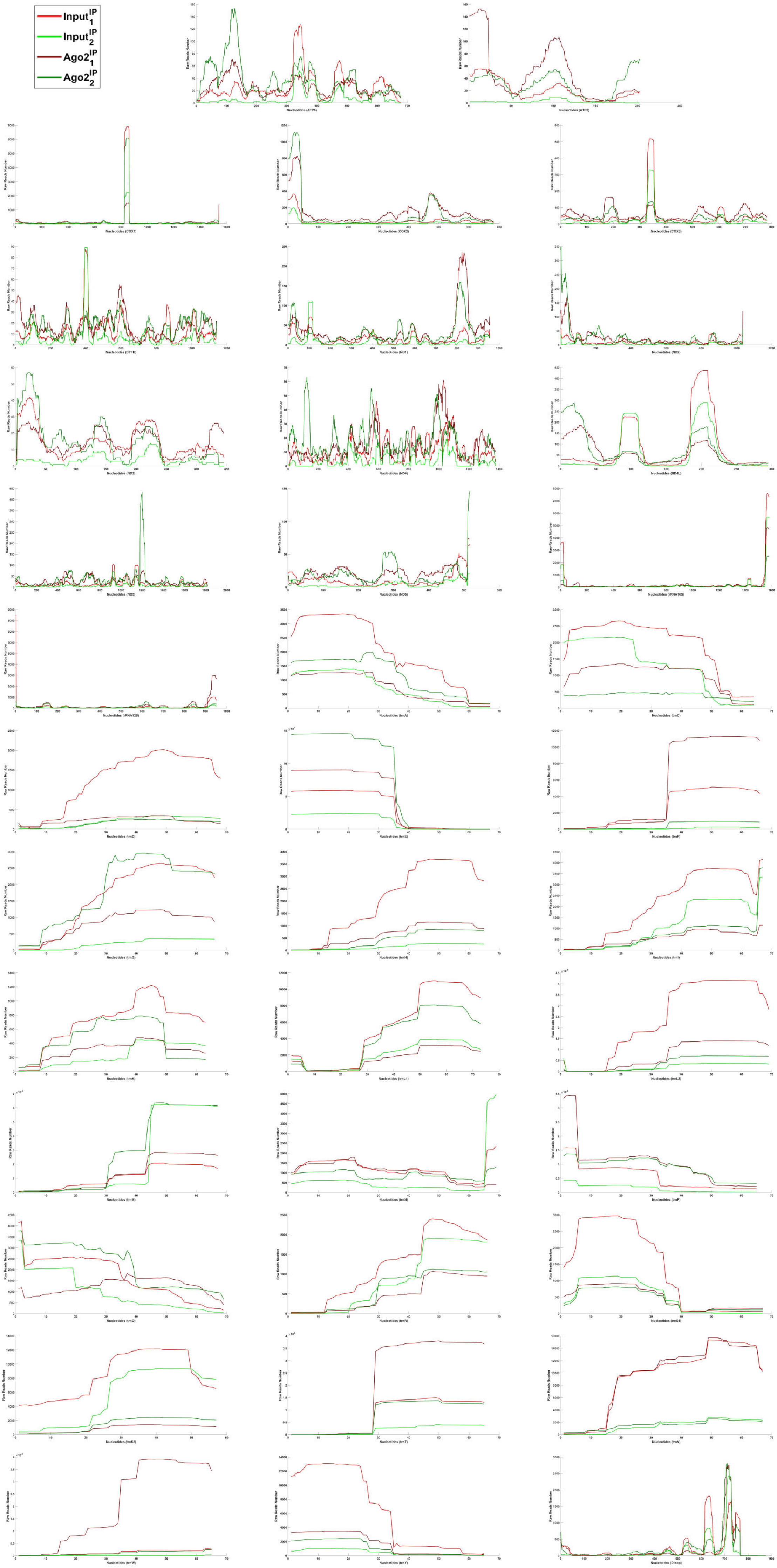
$Input_1^{IP}$
$Input_2^{IP}$
$Ago2_1^{IP}$
$Ago2_2^{IP}$
Raw Reads Number
Nucleotides (ATP6)
Nucleotides (ATP8)
Nucleotides (COX1)
Nucleotides (COX2)
Nucleotides (COX3)
Nucleotides (CYTB)
Nucleotides (ND1)
Nucleotides (ND2)
Nucleotides (ND3)
Nucleotides (ND4)
Nucleotides (ND4L)
Nucleotides (ND5)
Nucleotides (ND6)
Nucleotides (rRNA16S)
Nucleotides (rRNA12S)
Nucleotides (trnA)
Nucleotides (trnC)
Nucleotides (trnD)
Nucleotides (trnE)
Nucleotides (trnF)
Nucleotides (trnG)
Nucleotides (trnH)
Nucleotides (trnI)
Nucleotides (trnK)
Nucleotides (trnL1)
Nucleotides (trnL2)
Nucleotides (trnM)
Nucleotides (trnN)
Nucleotides (trnP)
Nucleotides (trnQ)
Nucleotides (trnR)
Nucleotides (trnS1)
Nucleotides (trnS2)
Nucleotides (trnT)
Nucleotides (trnV)
Nucleotides (trnW)
Nucleotides (trnY)
Nucleotides (Dloop)
