## Supplemental Figure 5 for "A new member in the Argonaute crew: the mt-miRNAs"

### Expression of small and long mitochondrial RNAs from kidney samples

**Figure S5** shows the expression of small and long mitochondrial RNAs across the entire mitochondrial genome in mouse.

In the following figure, we show that the small RNA transcriptional signatures at stages, day1 (P1), day3 (P3), and day7 (P7). There are three biological replicates for each stage, and for each biological replicate both long and small RNAs have been sequenced. Thus, there is one small and one long RNA sample for each biological replicate, providing the best comparison for matching transcriptional signatures. The samples are not specifically enriched for any small or long RNAs, thus they show a mix of coding and non-coding RNAs.

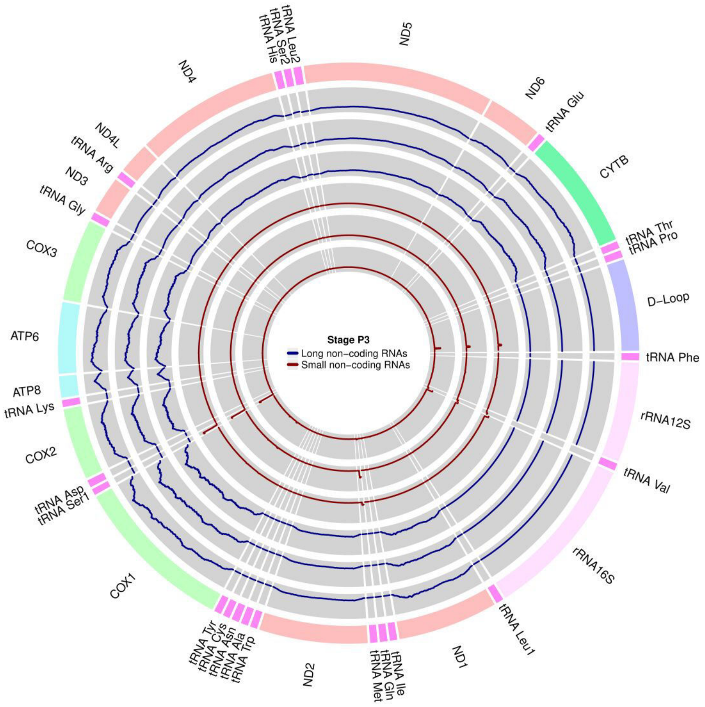
Stage P3
Long non-coding RNAs
Small non-coding RNAs
tRNA Leu2
tRNA Ser2
tRNA His
ND5
ND6
tRNA Glu
CYTB
tRNA Thr
tRNA Pro
D-Loop
tRNA Phe
rRNA12S
tRNA Val
rRNA16S
tRNA Leu1
ND1
tRNA Ile
tRNA Gln
tRNA Met
ND2
tRNA Tyr
tRNA Cys
tRNA Asn
tRNA Ala
tRNA Trp
COX1
tRNA Asp
tRNA Ser1
COX2
tRNA Lys
ATP8
ATP6
COX3
tRNA Gly
ND3
tRNA Arg
ND4L
ND4

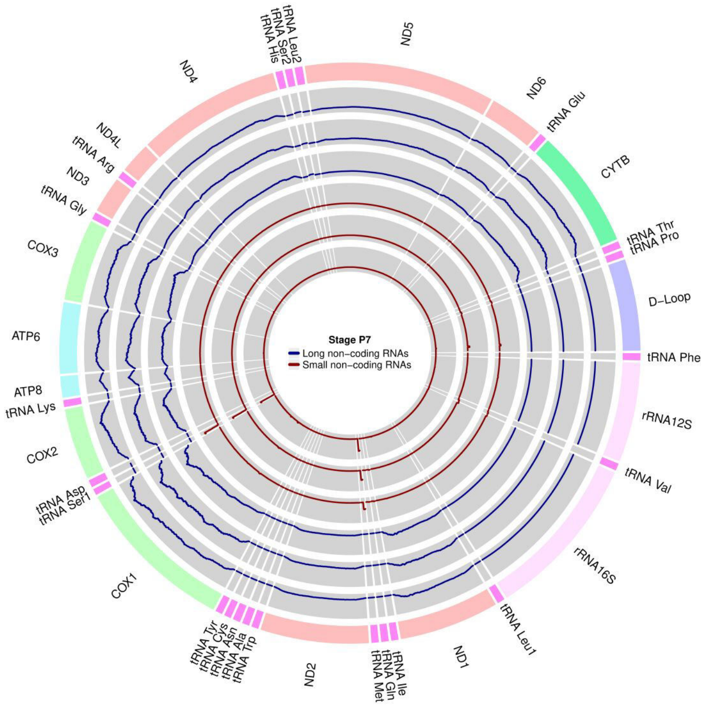
Stage P7
Long non-coding RNAs
Small non-coding RNAs
tRNA Leu2
tRNA Ser2
tRNA His
ND5
ND6
tRNA Glu
CYTB
tRNA Thr
tRNA Pro
D-Loop
tRNA Phe
rRNA12S
tRNA Val
rRNA16S
tRNA Leu1
ND1
tRNA Ile
tRNA Gln
tRNA Met
ND2
tRNA Tyr
tRNA Cys
tRNA Asn
tRNA Ala
tRNA Trp
COX1
tRNA Asp
tRNA Ser1
COX2
tRNA Lys
ATP8
ATP6
COX3
tRNA Gly
ND3
tRNA Arg
ND4L
ND4

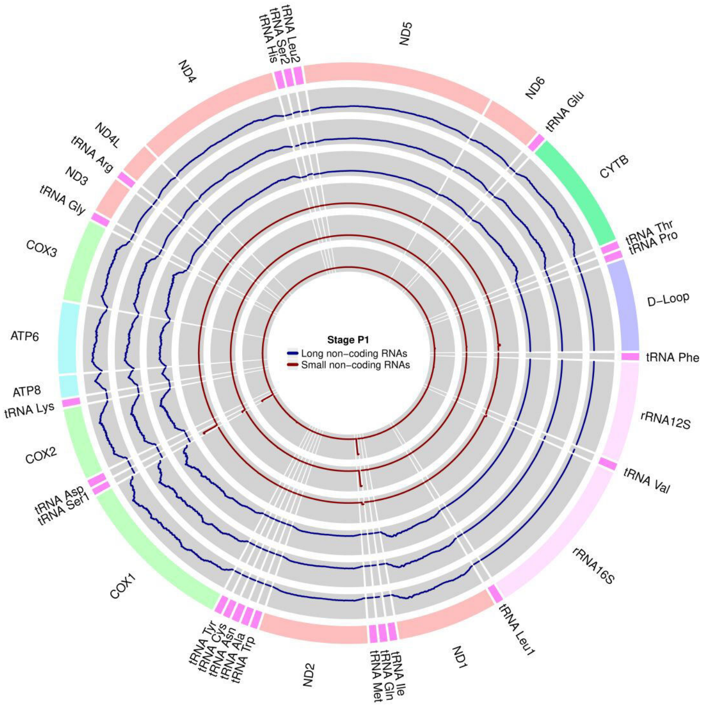
Stage P1
Long non-coding RNAs
Small non-coding RNAs
tRNA Leu2
tRNA Ser2
tRNA His
ND5
ND6
tRNA Glu
CYTB
tRNA Thr
tRNA Pro
D-Loop
tRNA Phe
rRNA12S
tRNA Val
rRNA16S
tRNA Leu1
ND1
tRNA Ile
tRNA Gln
tRNA Met
ND2
tRNA Tyr
tRNA Cys
tRNA Asn
tRNA Ala
tRNA Trp
COX1
tRNA Asp
tRNA Ser1
COX2
tRNA Lys
ATP8
ATP6
COX3
tRNA Gly
ND3
tRNA Arg
ND4L
ND4
