## Supplemental Figure 6 for "A new member in the Argonaute crew: the mt-miRNAs"

### CFLAR conservation in Primates

The **figure S6** shows the conservation of the CFLAR genomic region across five primates having a high-quality sequenced genome.

In the following figure, we show five dot-plots comparing the CFLAR genomic region in human (*Homo sapiens)* on the Y-axis to the respective region in chimpanzee (*Pan troglodytes*), bonobo (*Pan paniscus*), gorilla (*Gorilla gorilla*), orango (*Pongo abelii*), and olive baboon (*Papio anubis*) on the X-axis. In these plots each dot represents a sequence match between the two genomic regions, thus showing the conservation of the region on the diagonal of the plot. The region of the CFLAR 3’ UTR is highlighted in red, and the region of binding of mt-miRNA$_{Met}$ is highlighted in blue. The binding region is barely visible, as it has been misplaced to neighbor regions in all species, and it is not present on the diagonal.

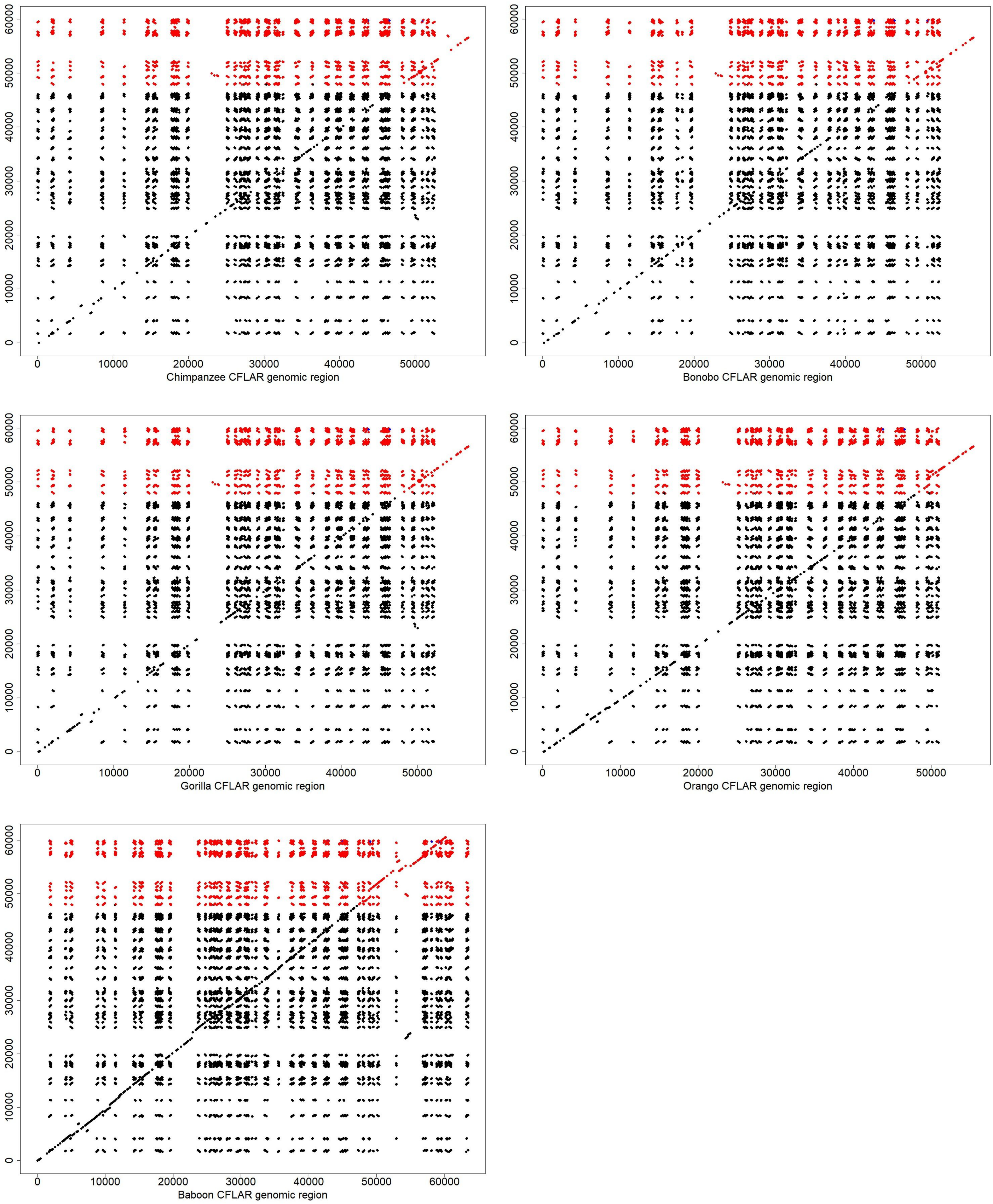
Chimpanzee CFLAR genomic region
Bonobo CFLAR genomic region
Gorilla CFLAR genomic region
Orango CFLAR genomic region
Baboon CFLAR genomic region
